# Elucidation of multiple high-resolution states of human MutSβ by cryo-EM reveals interplay between ATP/ADP binding and heteroduplex DNA recognition

**DOI:** 10.1101/2023.05.08.539930

**Authors:** Jung-Hoon Lee, Maren Thomsen, Herwin Daub, Gabriel Thieulin-Pardo, Stefan C. Steinbacher, Agnieszka Sztyler, Vinay Dahiya, Tobias Neudegger, Celia Dominguez, Ravi R. Iyer, Hilary A. Wilkinson, Edith Monteagudo, Nikolay V. Plotnikov, Dan P. Felsenfeld, Tasir S. Haque, Michael Finley, Julien Boudet, Thomas F. Vogt, Brinda C. Prasad

**Affiliations:** Proteros biostructures GmbH, Bunsenstr 7a, 82152 Martinsried, Germany; Current address: Roche Diagnostics GmbH, Nonnenwald 2, 82377 Penzberg, Germany; Current Address: Treeline Biosciences, 11180 Roselle St, San Diego, CA 92121, USA; Current Address: Script Biosciences, 2929 Arch Street, Philadelphia, PA 19104, USA; CHDI Management/CHDI Foundation, Princeton, NJ 08540, USA

**Author notes:** Contributed equally (Jung-Hoon Lee, Maren Thomsen). Correspondence: Brinda C. Prasad, PhD.

## Abstract

Human and mouse genetic studies have demonstrated a role for DNA mismatch repair (MMR) molecular machines in modulating the rate of somatic expansion of the huntingtin (*HTT)* CAG repeats, and onset and progression of Huntington’s Disease (HD). MutSβ, a key component of the MMR pathway, is a heterodimeric protein of MSH2 and MSH3 that recognizes and initiates the repair of extrahelical DNA extrusions. Loss-of-function of mouse *Msh3* and reduced-expression alleles of human *MSH3* lead to slower rates of somatic expansion and delayed disease onset in humans, signifying MSH3 as a promising therapeutic target for HD. Here we report biochemical and cryo-electron microscopy analyses of human MutSβ, demonstrating MutSβ undergoes conformational changes induced by nucleotide and DNA binding. We present multiple conformations of MutSβ including the DNA-free MutSβ compatible with homoduplex DNA binding, two distinct structures of MutSβ bound to (CAG)_2_ DNA, a sliding clamp form and a DNA-unbound, ATP-bound conformation. Along with evidence for novel conformational states adopted by MutSβ to initiate the MMR cascade, these structures provide a foundation for structure-guided drug discovery.

## Introduction

HD is a fatal neurodegenerative disorder caused by an expanded CAG repeat in the *HTT* gene. Individuals who inherit a *HTT* CAG-repeat length greater than 35 are at risk, and the disease is fully penetrant at CAG lengths greater than 39^1^. After the cloning of the *HTT* gene^2^, differential rates of CAG repeat somatic instability were observed in human CNS and peripheral tissues. Recent human genetic studies have shown that the number of uninterrupted *HTT* CAG triplet repeats, and not simply the number of glutamines dictate the timing of disease onset^3^. These studies in HD have revealed several genetic modifiers of the age onset and disease progression and remarkably, many of these genes encode proteins in the MMR pathway^4^. These findings intersect with prior reports implicating MMR genes in abrogating *Htt* CAG somatic expansion in mouse models of HD^5^. Both human and mouse genetics have identified *MSH3/Msh3*, encoding a subunit of the MutSβ heterodimeric complex, as a modifier of *HTT* CAG somatic expansion and disease onset.

The MMR pathway specifically identifies and corrects base-base mismatches and insertion/deletion loops (IDL) that form during DNA replication^6^. Defective aspects of MMR activity in humans contributes to cancers such as Lynch syndrome and constitutive MMR deficiency syndrome (CMMR-D)^7^. In contrast, the MMR pathway can be subverted to produce mutations and has been implicated in triplet repeat expansions that underlie several neurodegenerative diseases. Elevated amounts of MSH3 and the ensuing increase in repair activity hastens the onset of HD^4,8^. MutSα and MutSβ, eukaryotic homologues of the prokaryotic MutS, recognize mismatches and IDLs and initiate MMR by recruiting additional members of the pathway to execute the repair^9^. The MutSα heterodimer, composed of MSH2 and MSH6, shares functional similarities with the homodimeric bacterial MutS in its ability to detect one or two base pair mismatches. In contrast, MutSβ, composed of MSH2 and MSH3, recognizes IDLs ranging from 1 to 14 nucleotides and DNAs with 3’ single-stranded extensions^10-12^. Human MutSβ directly interacts with MutL endonucleases that introduce strand breaks, and Proliferating Cell Nuclear Antigen (PCNA), a DNA clamp and strand-directionality factor, to perform MMR^13^. MSH2 and MSH3 are paralogs and have similar overall domain organization. Previous crystallographic studies showed that MSH2 and MSH3 dimerize pseudo-symmetrically across their binding interface in a head-to-tail configuration^14^. The structures of both nucleotide binding sites are highly similar, containing Walker A and Walker B motifs which are evolutionarily conserved in all ATPases; the less conserved interunit signature loop projects from each subunit to the opposing nucleotide binding site^15-19^. In addition, mutagenesis and cryo-EM studies of *E. coli* MutS homodimer and human MutSα revealed conformational changes upon binding to different nucleotides and DNA^20-22^. Despite advances in our understanding of bacterial MutS and human MutSα/β structures, the exact mechanisms coupling MutSβ conformational changes to nucleotide exchange, DNA binding, and DNA release remain elusive. Here, we present biochemical and structural analysis of MutSβ in response to binding and release of nucleotides and DNA, suggesting a potential mechanism for MMR and its role in the somatic instability of *HTT* CAG repeats.

## Results

### MutSβ captured by cryo-EM reveals conformational transitions induced by binding to DNA substrates

To determine the structures of human MutSβ, we co-expressed two constructs encoding the full-length *MSH2* (NP_001393570.1) and *MSH3* (NP_002430.3) genes in insect cells and copurified them to homogeneity (Extended Data Fig. 1a-c). Size-Exclusion Chromatography with Multi-Angle Light Scattering (SEC-MALS) confirmed that the reconstituted complex is consistent with a calculated molecular mass of 232.1 kDa, corresponding to the MSH2-MSH3 heterodimeric complex. The catalytic activity of the purified full-length wild-type MutSβ was assessed by an *in vitro* ATPase assay (Extended Data Fig. 1d) and binding to DNA was demonstrated by Surface Plasmon Resonance (SPR) (Extended Data Fig. 1e). Cryo-EM analysis of vitrified specimens revealed different conformational states of particles, supported by clearly distinguishable three-dimensional (3D) maps (Fig. 1b-j).

**Fig. 1.**
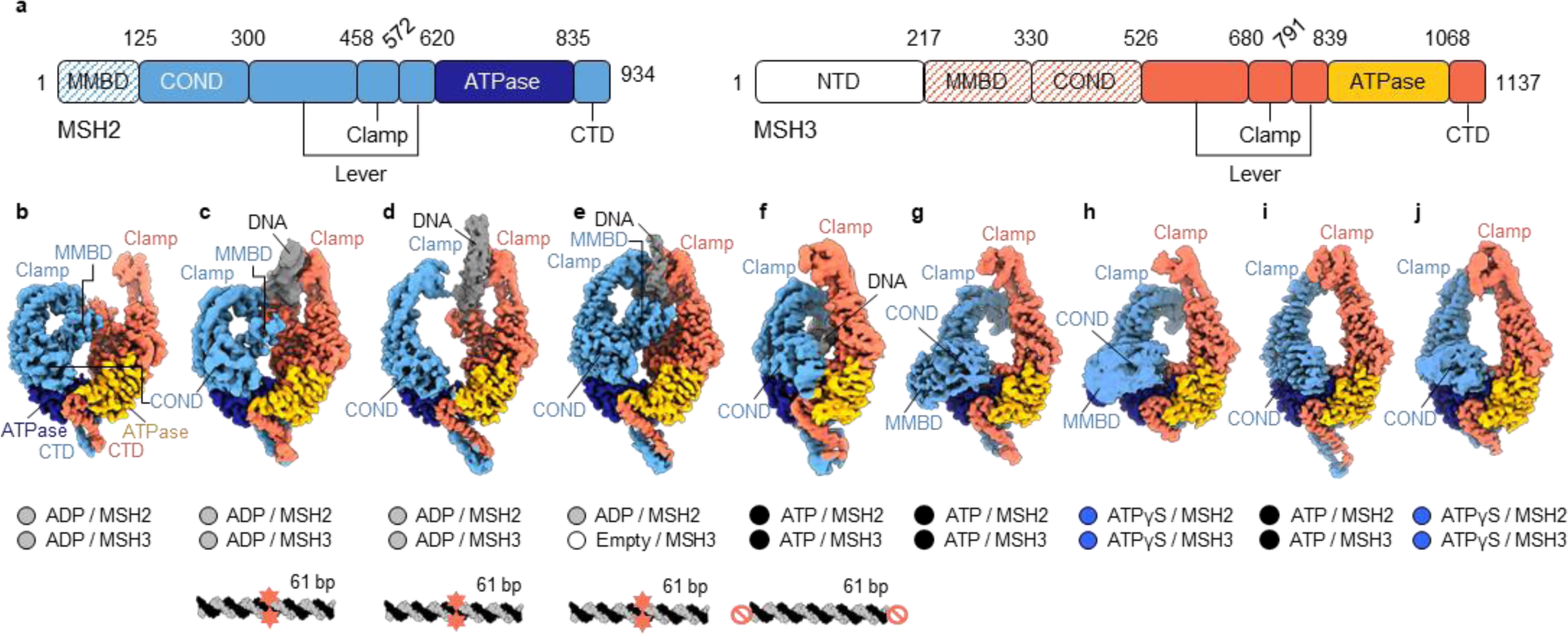
Human MutSβ heterodimer and its conformational heterogeneity captured by cryo-EM. **a,** Domain organization of human MutSβ subunits (mismatch-binding domain; MMBD, connector domain; COND, N-terminal domain; NTD, C-terminal domain; CTD). In all nine cryo-EM maps, the MSH3 NTD is not visible. The hatching pattern indicates the absence of density corresponding to the domains in several cryo-EM maps, presumably because of their flexibility (see results). **b-j,** Cryo-EM maps of nine different conformers of MutSβ (b, DNA-free open form, c, (CAG)_2_ DNA-bound open form with kinked MSH2 clamp, d, (CAG)_2_ DNA-bound open form, e, (CAG)_2_ DNA-bound canonical form, f, homoduplex DNA-bound sliding clamp, g and h, DNA-unbound form with kinked MSH2 clamp, i and j, DNA-unbound form with straight MSH2 clamp: See Table 1) The bound nucleotide at each subunit per conformer and the DNA substrate (mismatched bases with red stars and end-blocking of the DNA with red prohibition circles) used for cryo-EM are shown on the bottom panel. The same subunit color code is used throughout.

**Table 1.**
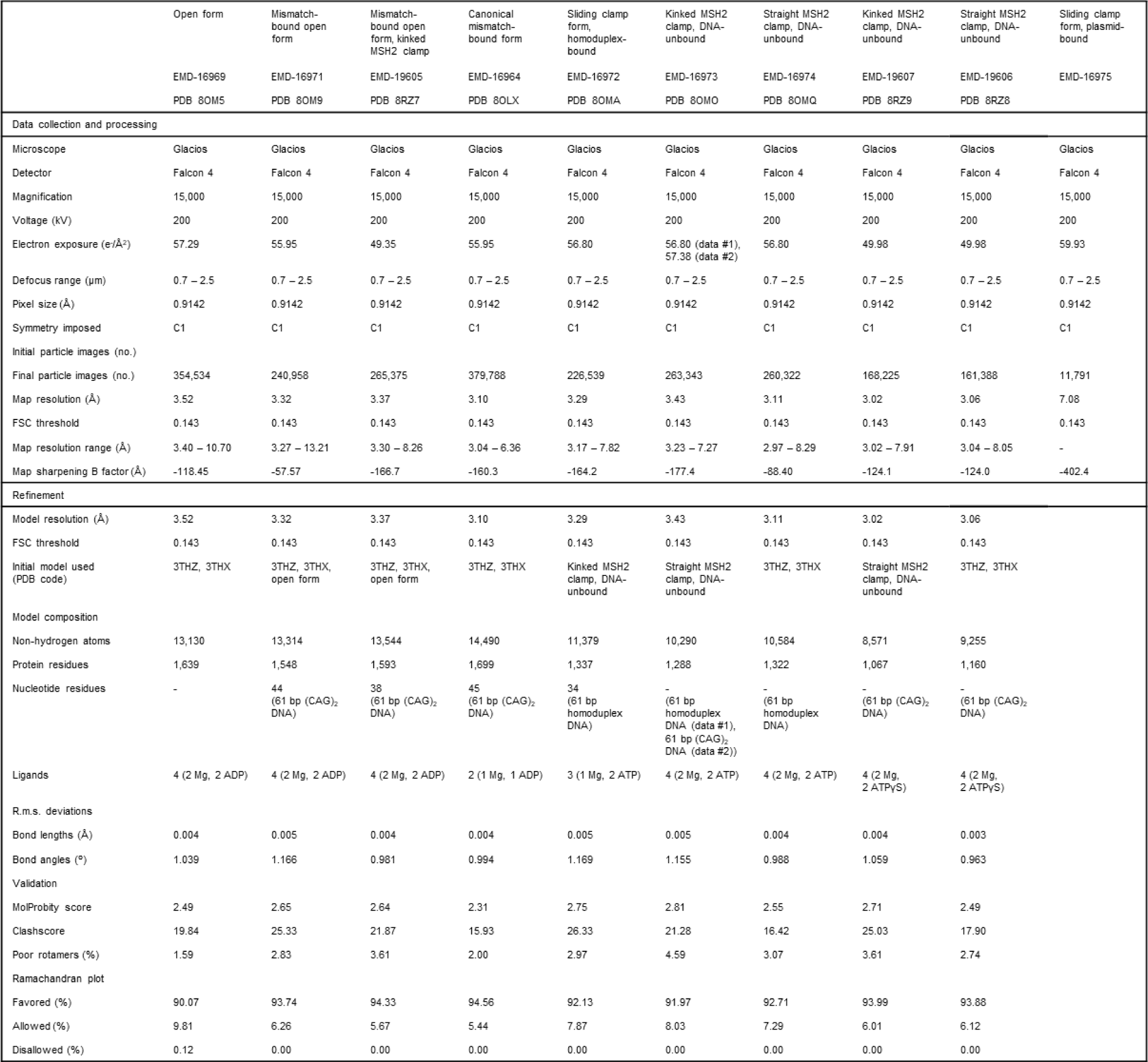
Cryo-EM data collection, refinement, and validation statistics.

The consensus reconstruction of DNA-free open form of MutSβ with 354,534 particles extended to an overall resolution of 3.52 Å as determined by gold-standard Fourier shell correlation (FSC; see Extended Data Fig. 2). The final map enabled us to unambiguously assign each domain of MSH2 and MSH3 to their corresponding electron densities (Fig. 1b). For model building, the crystal structures of the human MutSβ-DNA complex (PDB 3THZ and 3THX)^15^ were rigidly fitted into the cryo-EM map and then manually rebuilt according to the electron densities. The N-terminal domain (NTD) of MSH3 is not defined in the cryo-EM map, presumably because of its conformational flexibility. Intriguingly, MutSβ exhibited three different conformations in the presence of a 61 bp (CAG)_2_ loop containing DNA substrate. 2D classification and 3D reconstructions of particle images revealed two DNA-bound partially open forms of MutSβ at 3.37 Å and 3.32 Å that are distinct from a canonical mismatch-bound state at 3.10 Å (Fig. 1c-e). MSH2 had ADP bound in the canonical mismatch-bound form, while both MSH2 and MSH3 had ADP bound in the two open forms with the bound DNA. Three additional conformational states generated in the presence of homoduplex DNA were obtained from the EM specimen with ATP, showing more compact closed forms compared with those of MutSβ in more relaxed ADP-bound states (Fig. 1f,g,i, see below). Two additional conformers, bound to ATPγS at 3.02 Å and 3.06 Å, displayed compact closed forms highly reminiscent of the ATP-bound conformations reported above (Fig. 1h,j).

### DNA-free MutSβ adopts an open conformation with asymmetric clamps and ADP bound to both subunits

In the absence of DNA, MutSβ adopts a half-opened shape resembling a pretzel with the two subunits oppositely coupled at near-central domains and measures ∼148 Å in its longest dimension and ∼106 Å in width in the high-resolution cryo-EM map (Fig. 2a). The two subunits exhibit pseudo-twofold symmetry with a butterfly-shaped C-terminal ATPase heterodimer linked to lever and clamp domains at the opposite ends (see Fig. 1a for domain borders), an overall architecture resembling the bacterial MutS homodimer (Extended Data Fig. 2g). A reported *E. coli* apo MutS cryo-EM structure (PDB 7OU2)^21^ adopts a symmetric open conformation with a common dyad axis for the ATPase homodimer, with poorly resolved densities for the clamp domains. This conformation was also observed for *E. coli* MutS in the DNA scanning state (PDB 7AI5)^20^ and under DNA-free conditions for *N. gonorrhea* MutS (PDB ID 5X9W and 5YK4)^23^. By contrast, MSH2 in the open state of MutSβ is significantly bent, likely induced by interaction with the mismatch binding domain (MMBD) of MSH3, such that the MSH2 clamp is kinked inward by ∼95° toward MSH3. The straight conformation of the MSH3 clamp does not block the DNA entry channel of MutSβ compatible with DNA binding. The relative orientation of the domains of each subunit observed in the 3.52 Å cryo-EM structure resembles the previously determined crystal structures of the human MutSβ-DNA complex (PDB 3THZ)^14^ with an RMSD of 1.7 Å for 692 Cα atoms out of 934 MSH2 residues and 2.0 Å RMSD for 688 Cα atoms out of 1137 MSH3 residues. Additionally, we clearly observe the two clamps in this structure which were previously unresolved in the *E. coli* structure (PDB 7OU2). In the DNA-free open conformation, MSH2 and MSH3 subunits are tilted across each other by ∼42° relative to a canonical mismatch-bound structure, which we hypothesize is required for DNA binding (Extended Data Fig. 2h).

**Fig. 2.**
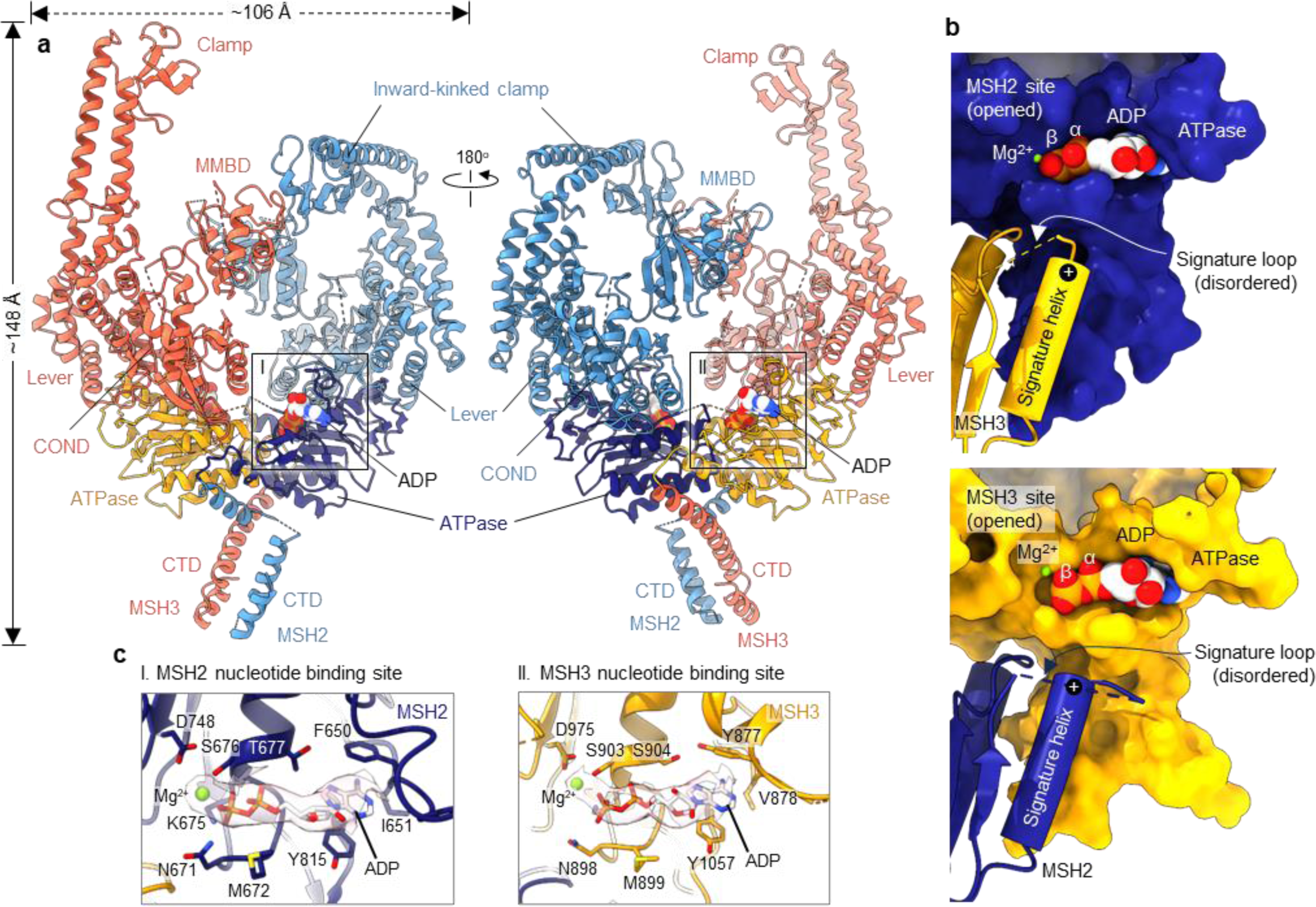
DNA-free MutSβ adopts an open, ADP-bound conformation with asymmetric clamps. **a,** Two views of a cartoon representation of MutSβ-ADP complex in the open conformation at 3.52 Å. **b,** Surface representation of MSH2 and MSH3 ATPase domains. In the open, ADP-bound conformer, the nucleotide is exposed without obstruction by the signature loop and helix from the opposing subunit. The positively charged end of the helix dipole is distanced from the phosphate groups of the bound nucleotide. **c,** Magnified views showing nucleotide binding sites of MutSβ in the open form. Electron densities for ADP-Mg are shown in pink (transparent).

Grid freezing of MutSβ in the absence of any DNA yields a complex with ADP-Mg bound to both subunits. This occurs even without adding ADP or ATP during protein purification, indicating that the ADP originates from ATP hydrolysis during protein expression (Fig. 2b,c). In our structures, ADP-Mg is solvent exposed at the α/β ATPase cores of MSH2 and MSH3, the signature loops are disordered (residues 714-721 and 940-949 of MSH2 and MSH3, respectively) and α-helices hosting the signature loops are located ∼11 Å farther away from the bound nucleotides within the active sites (Fig. 2b). Aligned with previous studies, we found that the interface between ADP-Mg and MSH3 involves conserved aromatic, hydrophobic, and charged residues. MSH3 Walker motifs^15^ coordinate the ADP phosphate groups and magnesium. Nucleotide binding pockets are essentially the same for MSH2 and MSH3 (Fig. 2c).

### Limited contacts with DNA orient MutSβ in two different half-open states

We determined three distinct conformational states of full-length MutSβ bound to mismatched DNA using cryo-EM. The first conformation was the canonical mismatch-bound state of the 3.10 Å cryo-EM structure resembling published crystal structures of truncated MutSβ-DNA complexes^14^ (Fig. 3a,b and Extended Data Fig. 3l). The MSH2-MSH3 heterodimer adopts the typical mismatch-bound state, with the ATPase domain and C-terminal domain (CTD) forming an asymmetrical dimeric structure while the clamp and lever domains extend out. The DNA interacts with MMBD of MSH3 and partially with its counterpart in MSH2, as seen in previous crystal structures^14,24,25^ (Fig. 3c). In our cryo-EM structure, the DNA is sharply kinked by ∼67° at the IDL of (CAG)_2_ mis-paired nucleotides. The two conserved residues, tyrosine 254 and lysine 255 of MSH3, penetrate the DNA at the IDL to stretch apart the DNA strands (Fig. 3c and Extended Data Fig. 3m). At position 42 in MSH2, phenylalanine engages in a π stack with the fourth base of the (CAG)_2_ insertion, while at position 254 in MSH3, the tyrosine participates in a π-π stacking interaction with the unpaired base in the complementary DNA strand (Fig. 3c). However, a portion of this DNA loop is disordered (Fig. 3a,c). Consistent with previous crystallography studies^14^, ADP was bound to the MSH2 site, even though ADP or ATP was not intentionally included in the sample. However, no density corresponding to a nucleotide was detected in the MSH3 catalytic pocket (Fig. 3b,d). During examination of 2D class averages, the second conformation was observed in a subset of MutSβ-DNA complexes adopting a distinct open conformation of MSH2. Using this subset of particles, we solved a new structure of the heteroduplex DNA-bound MutSβ complex by cryo-EM distinct from the canonical DNA mismatch-bound structure described in Figure 3. A 3D reconstruction was calculated based on extensive 2D classification of cryo-EM data interpreted as the MSH2 and MSH3 subunits in complex with a 61 bp (CAG)_2_ loop DNA (Extended Data Fig. 3a,g-k). In this complex, the MutSβ-DNA interaction is mediated by multiple contacts between DNA and the MMBD and clamp domains of MSH3, exhibiting substantial movement of MSH3 clamp and MMBD domains by ∼7 Å and ∼5 Å, respectively, as compared to the DNA-free structure (Fig. 4a,c and Extended Data Fig. 3n). Despite not being involved in DNA binding, the MSH2 clamp moves outward by ∼27 Å relative to its DNA-free open form when bound to the heteroduplex DNA. The cryo-EM structure reveals that the DNA undergoes a sharp kink of ∼73° at the IDL of (CAG)_2_ mis-paired nucleotides (Extended Data Fig. 3o). Intriguingly, only the MSH3 subunit of the MutSβ heterodimer is involved in DNA binding (Fig. 4a,c). These DNA contacts with the MMBD of MSH3 are similar to those of the canonical mismatch-bound state of MutSβ, while MSH3 clamp interacts differently with DNA, thus representing a partially open conformation of MutSβ assembled onto the heteroduplex DNA (Fig. 4a,c and Extended Data Fig. 3p). We observed that both the MSH2 and MSH3 ATPase nucleotide binding pocket are occupied by ADP-Mg (Fig. 4e).

**Fig. 3.**
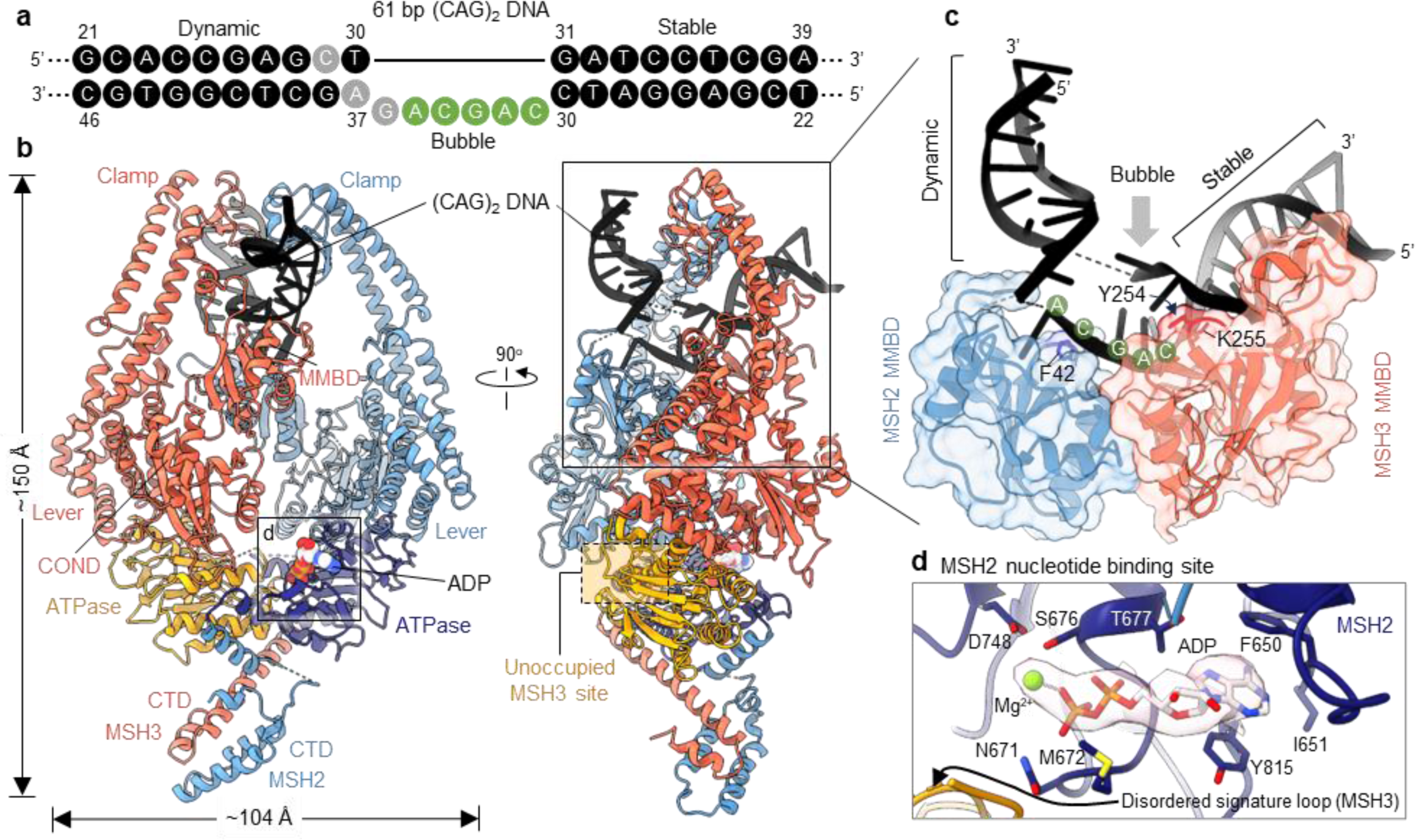
MutSβ bound to (CAG)_2_ DNA in a canonical mismatch-bound conformation. **a,** Nucleotides are represented as solid circles in black (modelled) or grey (disordered). Green circles indicate the modelled mis-paired bases of two CAG repeats. **b,** Two orthogonal views of a graphic representation of the MutSβ-ADP complex bound to (CAG)_2_ mismatched DNA. **c,** A magnified side view of the DNA and two MMBDs. Three key residues, MSH2 F42, and MSH3 Y254 and K255, are shown as sticks. **d,** Close-up view of the MSH3 nucleotide binding pocket of MutSβ in the mismatch-bound state. Electron density for ADP-Mg is shown with transparent shell.

**Fig. 4.**
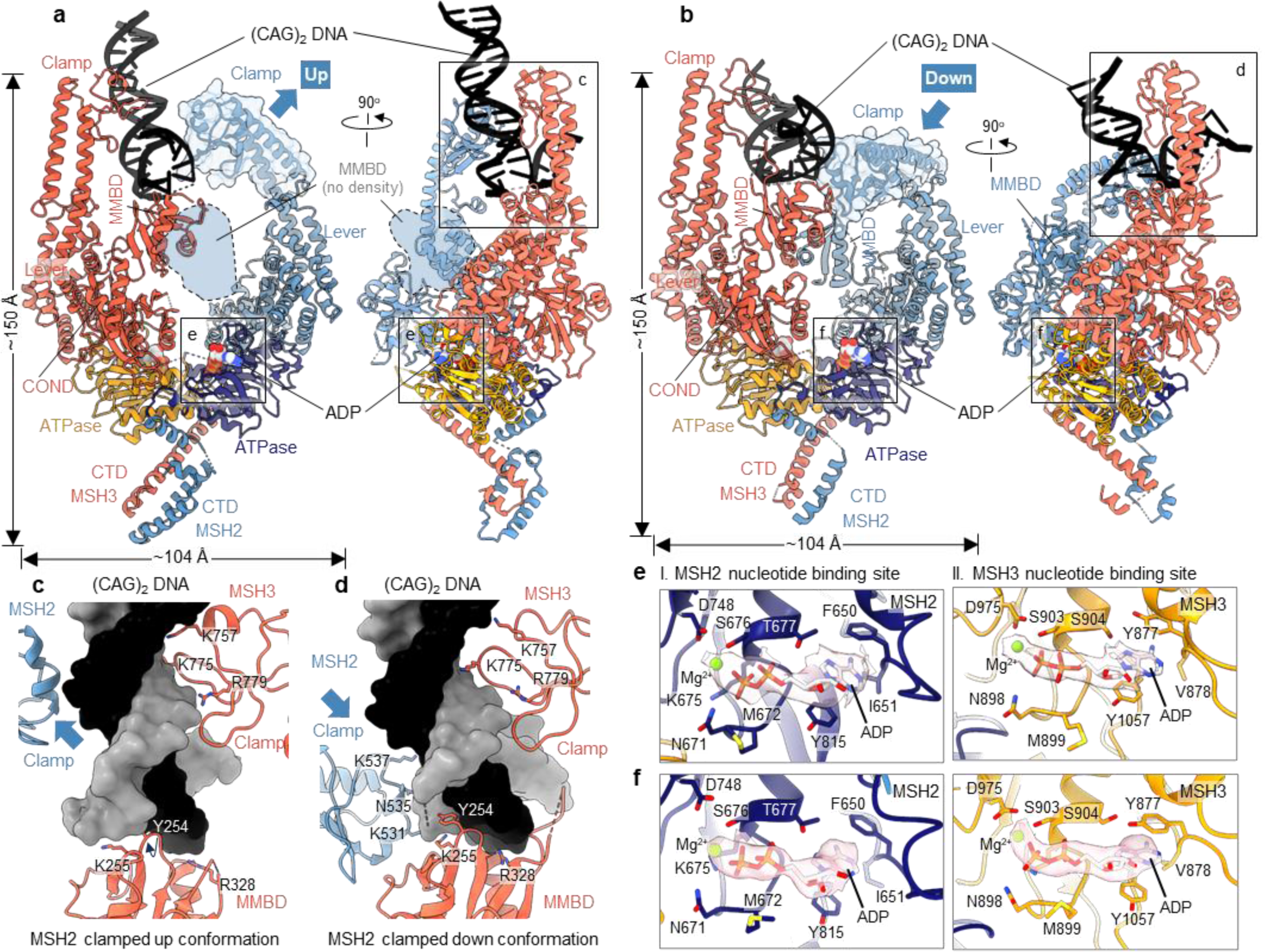
Limited contacts with the mismatched DNA orient MutSβ in the open states. Ribbon diagrams showing two views of the MutSβ-ADP complex with MSH2 clamped up (a) and down (b) in the (CAG)_2_ DNA-bound open conformations at 3.32 Å (PDB 8OM9) and 3.37 Å (PDB 8RZ7), respectively. Close-up views of the interfaces between MutSβ and DNA in the MSH2 clamped up (c) and down (d) conformations. Magnified views showing nucleotide binding sites of MutSβ in the DNA-bound open form with MSH2 clamped up (e) and down (f) conformations. Electron densities for ADP-Mg are shown with a transparent shell (pink).

The third, alternative conformer of the MutSβ complex, bound to the same heteroduplex DNA, exhibited a distinct open conformation and was obtained through cryo-EM when incubating the MutSβ-DNA complex in the presence of AMP-PNP (Fig. 4b). The primary population of particles reconstructed in the 3.37 Å cryo-EM map revealed a MutSβ conformer almost identical to the one observed in the DNA-free ADP-bound structure, with RMSD of 0.933 Å for 765 Cα atoms in MSH2 (∼82%) and 1.171 Å RMSD for 797 Cα atoms in MSH3 (∼70%) (Extended Data Fig. 4a,g). The 61 bp (CAG)_2_ loop DNA occupies a near-peripheral area of MutSβ formed by the MSH2 and MSH3 clamps (Fig. 4b). While the MSH3 clamp and MMBD exhibit similar involvement in DNA binding compared to the structure shown in Fig. 4a, the MSH2 clamp undergoes an inward movement by 95° toward the DNA, resulting in a distinct binding mode in the alternative open conformation of MutSβ assembled to the mismatched DNA (Fig. 4d and Extended Data Fig. 4h). Despite an approximately 285-fold molar excess of AMP-PNP added to the preformed MutSβ in complex with DNA, our cryo-EM analysis indicated that most protein-DNA complexes maintained their ADP-bound state in both nucleotide-binding pockets of the MSH2 and MSH3 subunits (Fig. 4f and Extended Data Fig. 4a). Observation of these previously uncharacterized MutSβ conformations exhibiting weak association with IDL-DNA is consistent with the recently proposed functional model and biophysical experiments suggesting an induced-fit (2-step binding) recognition of mismatch by MutSα^22^.

### MutSβ undergoes an ATP-dependent conformational change towards a compact structure on DNA

A correlation between the ATP binding of MutS proteins and their structural change has been previously suggested^26-30^. Upon binding to heteroduplex DNA, MutS rapidly exchanges ADP for ATP, and is proposed to undergo a conformational change to form a sliding clamp^30^. Recent cryo-EM and biochemical studies of the bacterial MutS have revealed this conformational shift, adopting a compact structure in the presence of ATP, AMP-PNP, or ADP vanadate^20,21^.

To test whether ATP binding contributes to MutSβ’s ability to undergo conformational changes while binding to DNA, we conducted cryo-EM measurements using the full-length MutSβ mixed with a ∼1.8 kb linearized homoduplex plasmid DNA in the presence of ATP (Fig. 5a-c). Because the long DNA strands stretched across the vitrified holes on the cryo-EM grid, MutSβ displayed a preferential orientation on the DNA. The DNA-bound form, characterized by the highly recognizable electron density of the DNA, generated well-resolved 2D class averages, and the 3D reconstruction produced a 7.08 Å cryo-EM map (Fig. 5c) which revealed a structure distinct from the canonical mismatch bound state. Despite the low resolution, we observed that the binding of MutSβ to long homoduplex plasmid DNA with ATP results in the tilting of its two subunits and bending of the MSH2 clamp, which pushes the DNA downwards in relation to its mismatch-bound conformation and positions it in the central pore of MutSβ (Extended Data Fig. 5). Using a shorter 61bp end-blocked homoduplex DNA, we determined a 3.29 Å cryo-EM structure of MutSβ upon ATP binding (Fig. 5d and Extended Data Fig. 6a,b-f). The consensus reconstruction showed a clear density for MutSβ and the bound ATP, although the DNA was poorly resolved. In contrast to the heteroduplex DNA-bound proteins, the MMBDs of the two subunits and the MSH3 connector are not defined in the cryo-EM map, presumably due to their conformational flexibility. MSH2 had a clear density for its bent clamp towards DNA, positioning the DNA in the central pore of the MutSβ heterodimer. The 3D refinement indicated a substantial conformational flexibility of the MSH2 connector domain, with its relative position rotating downward by ∼180°, while the mismatch-binding domain was not observed (Extended Data Fig. 6q). The map displayed a continuous density of DNA extending from the MSH2 connector to its bent clamp and is modeled over a total of 17 bp. Nevertheless, we observed a lower resolution for the DNA compared to the rest of the cryo-EM map, likely due to the inherent flexibility and heterogeneity in the position of the DNA. Despite the limitations of the DNA density, the ATP-bound MutSβ appears to bind to DNA primarily through positively charged clusters partly embedded in MSH2 connector, lever, and clamp domains (Fig. 5e). Our two DNA-bound sliding clamp models of MutSβ demonstrate that the MSH2 connector can adopt inward or outward conformations (Fig. 5f). This movement does not appear to be directly influenced by the position of the DNA, although the exact cause of the MSH2 connector domain movement in the two different sliding clamp forms remains unclear. In one model, based on the map of MutSβ bound to a long homoduplex plasmid DNA, the MSH2 connector is positioned ∼27 Å away from the DNA. In the other model, based on the map of MutSβ bound to a short 61 bp homoduplex DNA, the MSH2 connector domain appears to move inward toward one end of the DNA, suggesting flexible movement. As expected, the observed ATP-bound MutSβ-DNA complex highly resembles the previously determined cryo-EM structure of *E. coli* MutS sliding clamp (PDB 7AIC; see Extended Data Fig. 6r)^20^; the sliding clamps display similar overall shapes and domain rearrangements, highlighting the structural similarities between human and bacterial proteins. Notably, a specific subunit’s clamp domain (e.g., MSH2 of human MutSβ and one MutS of *E. coli* MutS homodimer characterized by the kinked clamp in both structures, the RMSD (Cα atoms) is ∼2.95 Å) is observed to be sharply kinked towards DNA at the interface between the lever and clamp domain in both cryo-EM structures. Additionally, we observed that the tilting of ATPase domains in the MutSβ sliding clamp form upon ATP binding also occurs in a manner similar to that found in *E. coli* MutS (Fig. 5h)^20,21^. The closing motion of the ATPase domains in the MutSβ sliding clamp is associated with the signature helices, where the amino terminus of each helix interacts with the γ-phosphate of the bound ATP in the opposite subunit. This interaction leads to the closure of both the MSH2 and MSH3 sites, resulting in well-ordered signature loops and a more compact structure compared to the relatively relaxed canonical mismatch-bound state (Fig. 5g).

**Fig. 5.**
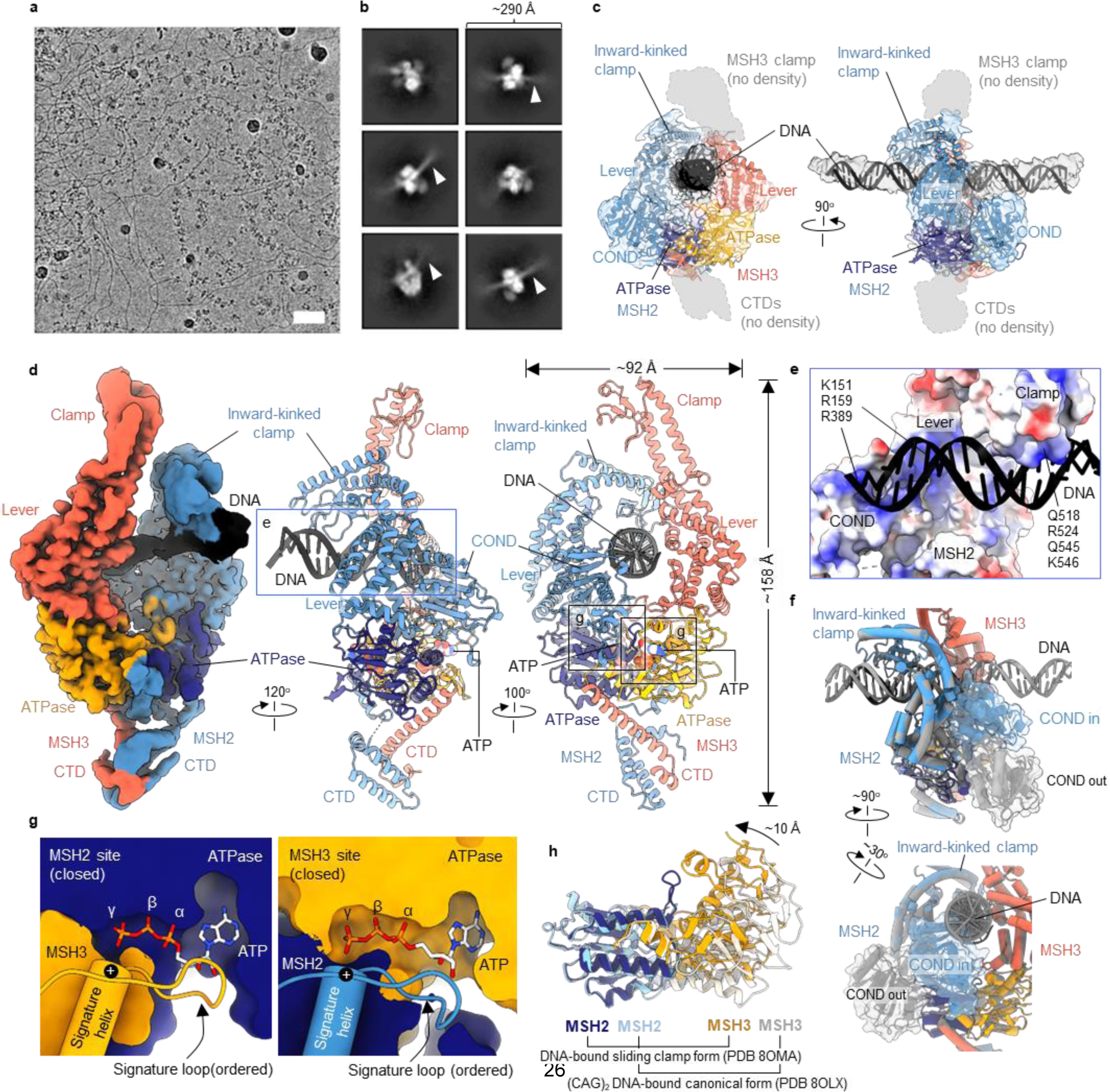
MutSβ undergoes ATP-dependent conformational change towards sliding clamp on DNA. **a,** Representative raw particles from an original micrograph (scale bar, 37.3 nm). **b,** Representative reference-free 2D class averages of MutSβ-ATP complex bound to a 1.8 kb homoduplex plasmid. Arrows point towards the DNA. **c,** Model of MutSβ-ATP-DNA complex fit into a 7.08 Å cryo-EM map. **d,** Three views of cryo-EM reconstruction of MutSβ bound to a 61 bp homoduplex DNA at a resolution of 3.29 Å (PDB 8OMA) (far left) and ribbon diagram of the structure. **e**, A close-up view of the interface between MSH2 and bound DNA. Key residues of MSH2 involved in DNA binding are highlighted in boxes. **f,** Two images showing the overlay of two structures of MutSβ-DNA-ATP complexes (homoduplex plasmid-bound sliding clamp in grey, 61 bp homoduplex-bound sliding clamp in color). **g,** Slice through surfaces displaying MSH2 (left) and MSH3 (right) nucleotide binding pockets of the 61 bp homoduplex DNA-bound MutSβ-ATP complex. The signature loops and helices from opposite subunits are shown as ribbons, and the bound ATPs are shown as a stick model. **h,** Ribbon diagrams of the canonical and sliding clamp forms of MutSβ bound to DNA, based on superposition, showing the tilting of the ATPase domains in the sliding clamp form.

To investigate in detail the role of ATP in conformational transitions of MutSβ-DNA complex, intra-molecular FRET experiments were performed using an end blocked 70bp (CAG)_2_ DNA labeled with a donor and acceptor dye on opposite strands of double stranded DNA stretches. The hypothetical FRET model in absence and presence of MutSβ is shown (Extended Data Fig.7a). An increase in the FRET efficiency of the labeled DNA was observed upon addition of MutSβ only in the case of the heteroduplex DNA and not with 70bp control homoduplex DNA (Extended Data Fig.7b). Based on our FRET model, DNA bending induced by MutSβ binding to the (CAG)_2_ DNA increases FRET between the donor and acceptor dyes. The inherent FRET efficiency of the (CAG)_2_ DNA was observed to be higher compared to the homoduplex DNA (Extended Data Fig.7b), possibly because of intrinsic flexibility of the (CAG)_2_ DNA^31^. Consequently, the difference in FRET efficiency of (CAG)_2_ DNA with and without MutSβ was smaller, but statistically significant (Extended Data Fig.7b). We incorporated other 70bp IDLs (CA, (CAG)_1_ and (CAG)_4_) in our FRET assay and found that these heteroduplex DNAs also resulted in a significant increase in the FRET efficiency in presence of MutSβ (Extended Data Fig.7b), consistent with the model of DNA kinking upon MutSβ binding. Interestingly, the addition of ATP together with MutSβ led to a decrease in the FRET efficiency of MutSβ-bound DNA (Extended Data Fig.7b) suggesting that ATP releases MutSβ from the loop out by sliding away or by releasing MutSβ altogether from the DNA. The three states (DNA-unbound MutSβ, DNA-bound MutSβ, and DNA-bound MutSβ in presence of ATP) with respect to the FRET efficiencies are clearly discernible in the heat map (Extended Data Fig.7c).

### Rearrangement of MutSβ clamp resembles bacterial MutS sliding clamp structures

The 2D class images of the MutSβ-61 bp homoduplex DNA-ATP complex in its sliding clamp form revealed a subset of particles that were not bound to DNA (Fig. 6a,c and Extended Data Fig. 6a). The pseudo two-fold symmetry between MSH2 and MSH3, as evident in their domains including the clamps, was analyzed. Two structures of the DNA-unbound MutSβ with ATP bound in both subunits were obtained, each with a different conformation of the MSH2 clamp (Extended Data Figs. 6g-p). The first is the 3.11 Å cryo-EM structure that shows straight clamps of both subunits bound to ATP-Mg without DNA (Fig. 6a). The second structure of the DNA-unbound form, at 3.43 Å, reveals that the overall architecture of MutSβ, including the bent clamp of MSH2, is nearly identical to the DNA-bound sliding clamp state of the MutSβ-ATP complex (Fig. 6c,e). An extra density adjacent to the MSH2 connector, corresponding to the N-terminal MMBD was identified, but the domain structure was not built due to the limited quality of the density (Extended Data Fig. 6p). The electron densities of the N-terminal, mismatch, and connector domains of MSH3 were not observed, presumably due to their flexibility. In both forms without DNA, the ATPase domains of both subunits are closed by a well-ordered signature loop and helix from the opposite subunit, and ATP-Mg is present in the nucleotide-binding pockets of MSH2 and MSH3 (Fig. 6f and Extended Data Fig. 8a).

**Fig. 6.**
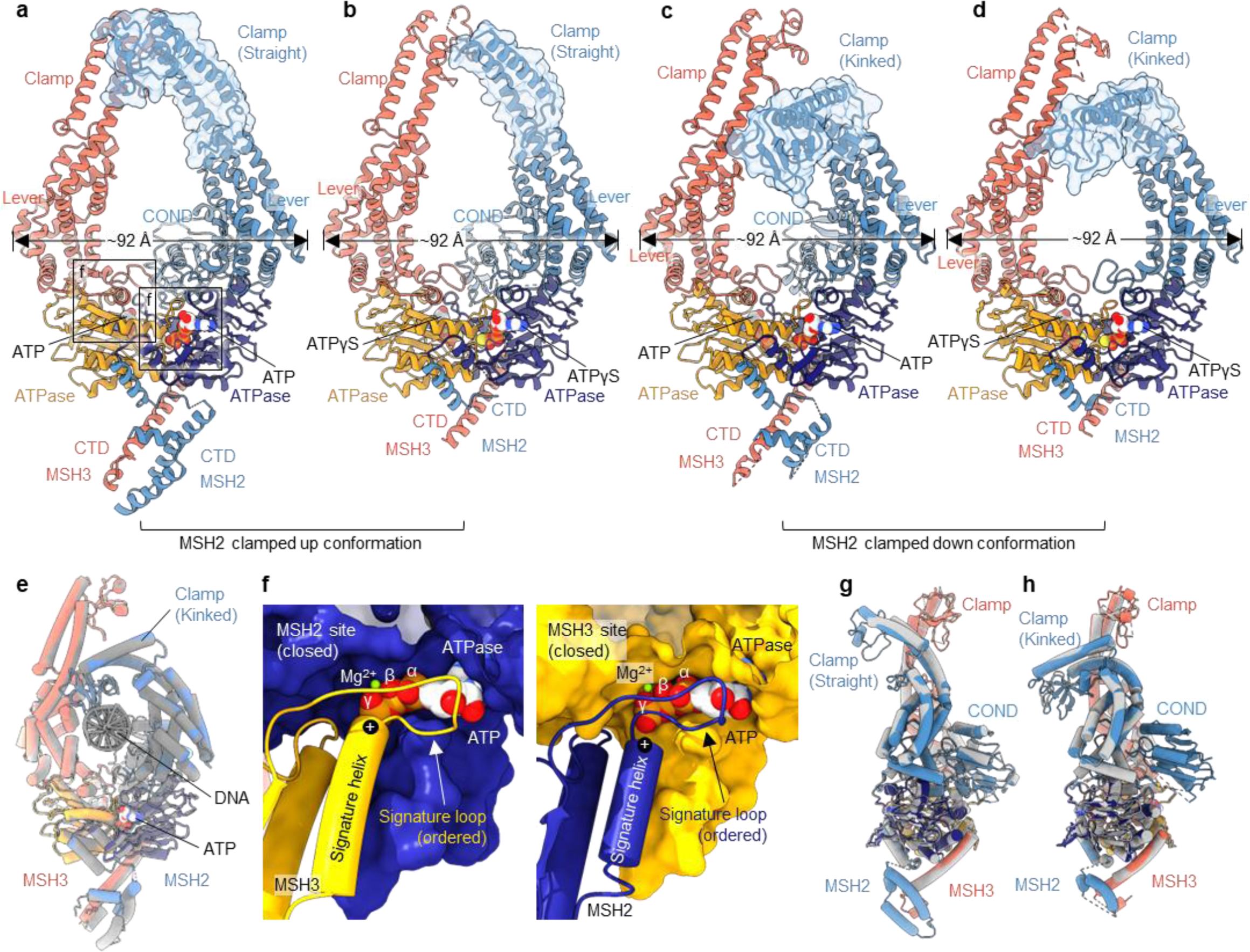
Cryo-EM structures of DNA-unbound MutSβ-ATP and MutSβ-ATPγS complexes. a-d, Cartoon representation of the MutSβ-ATP complex with a straight (a) or kinked (c) MSH2 clamp, and the MutSβ-ATPγS complex with a straight (b) or kinked (d) MSH2 clamp. Different conformations of the MSH2 clamp in the four structures are shown with transparent surfaces in blue overlaid with ribbon cartoons. **e,** Superposition of the two MutSβ-ATP complexes in the absence and presence of 61 bp homoduplex DNA. **f,** Close-up views showing nucleotide binding pockets of MutSβ in the DNA-unbound MutSβ-ATP complex with a straight MSH2 clamp. Overlays of each conformer of MutSβ-ATP in color with MutSβ-ATPγS in grey (g, straight MSH2 clamp form; h, kinked MSH2 clamp form).

We also assessed the impact of ATP analogs on conformational changes in the MutSβ-DNA complex by cryo-EM. The addition of AMP-PNP to MutSβ in complex with (CAG)_2_ DNA did not effectively induce conformational changes toward the sliding clamp form (Extended Data Figs. 4 and 9l). However, the addition of ATPγS to the preformed MutSβ-DNA complex resulted in the effective release of the bound DNA from MutSβ, leading to the determination of two cryo-EM structures depicting DNA-unbound MutSβ with ATPγS bound to both the MSH2 and MSH3 subunits (Fig. 6b,d and Extended Data Fig. 9a-k). These observations were further validated by additional FRET and MST assays where a significantly diminished efficacy in releasing the bound DNA from MutSβ upon addition of AMP-PNP, relative to ATP or ATPγS was observed (Extended Data Fig. 9l), suggesting a lower binding affinity of MutSβ to AMP-PNP compared to ATP or ATPγS^9^. A structural comparison between MutSβ-ATP and MutSβ-ATPγS complexes, featuring either a straight or bent MSH2 clamp, reveals no other significant differences in their conformation (Fig. 6g,h and Extended Data Fig. 9m,n). This finding supports the notion that ATP or ATPγS binding results in conformational rearrangement of MutSβ towards its compact sliding clamp form, underscoring the importance of nucleotide binding in modulating MutSβ conformation. Intriguingly, a comparison of the two DNA-unbound MutSβ-ATP complex structures with bacterial MutS shows that a similar conformational change of clamp domain was observed in the DNA-bound sliding clamp structures of *E. coli* MutS in the presence of ATP analog. This suggests a general mechanism for the conformational change in the sliding clamp state of MutS family proteins (Extended Data Fig. 8b). Comparison to DNA-free conformations of *E. coli* MutS with AMP-PNP in both subunits to the DNA-unbound MutSβ loaded with ATP in both subunits reveal similarity, confirming a conserved mechanism of action for all MutS molecular machines^21,32^.

## Discussion

We present nine different high-resolution cryo-EM maps of the heterodimeric full-length human MutSβ, the most comprehensive conformational landscape reported to date of this critical MMR protein complex. Several structures show significant similarity to structures of the *E. coli* MutS homodimer. For example, our 3.52 Å DNA-free open form of MutSβ demonstrates a similar pseudo-twofold symmetry to a butterfly-shaped C-terminal ATPase heterodimer shared with the reported *E. coli* MutS cryo-EM structure^20,21^. However, we also resolved the two clamp domains of MutSβ, revealing an asymmetry between the kinked MSH2 and extended MSH3 domains not clearly observed in the previous MutS cryo-EM structure (Extended Data Fig. 2). In another example, our 3.11 Å DNA-unbound form of MutSβ previously exposed to homoduplex DNA and ATP shows a striking similarity to mismatched-bound MutS sliding clamp, particularly in the angle adopted between the clamp and lever domains of MSH2 and one of the MutS subunits (Extended Data Fig. 8b). Although our 3.10 Å cryo-EM structure of MutSβ bound to a (CAG)_2_ IDL resembles the canonical mismatched DNA bound state as already known from crystal structures using the truncated MutSβ-DNA complexes^14^ (Extended Data Fig. 3), we also identified a novel 3.32 Å DNA-bound open conformation (Fig. 4). Differences between these two conformations highlight the potential for significant domain movements, such as the >10 Å inward movement of the MSH2 and MSH3 clamp domains towards the DNA, required to achieve the DNA-bound canonical form (Extended Data Fig. 3p). Our study provides a more complete understanding of MutSβ function in the initiation of the MMR cascade. We observed that MutSβ either slides off DNA or is biased by ATP to adopt, at least for some time, the double ATP bound, DNA-unbound state similar to the sliding clamp state^20^. We also observed a conformation corresponding to the double ADP-bound state free in solution. ADP-bound MutSβ spontaneously binds and scans homoduplex DNA; upon encountering an IDL-loop, it forms the mismatch-bound state (reported Kd ∼1-0.1 nM)^14^. The release from the IDL loop and conformational change into the sliding clamp depends on ATP binding to both MSH2 and MSH3. It suggests that MSH3 (and MSH2) hydrolyzes ATP following release from DNA or that the hydrolysis initiates clamp opening of MutSβ and hence the transition to the open conformation. Alternatively, we cannot preclude the possibility that MutL complex binding facilitates the ATP hydrolysis in MutSβ while being in the sliding clamp state, resulting in a quicker reset of MutSβ after having successfully initiated the MMR pathway. In either case, we acknowledge the requirement for a closed catalytic cycle where the ejection of γ-phosphate is required for MutSβ complex release from DNA and thus render the molecular complex competent to bind to DNA again. We demonstrate that cycle and key conformations of MutSβ are evolutionarily conserved as they have clear counterparts in the previously described *E. coli* cycle^20,21,32^. ATP consumption is coupled with IDL recognition, as MutSβ seems to be effectively inhibited by ADP in MSH2 until this inhibition is lifted by a chain of events leading to ADP-ATP exchange at both nucleotide binding sites in the sliding clamp state^33^.

Despite the additional insights of the conformational heterogeneity of the MutSβ complex across both DNA-free and DNA-bound forms, we still do not fully understand the relationship between the different conformations and nucleotide binding or ATP hydrolysis. The previous crystallographic study of MutSβ investigated a truncated human MutSβ bound to IDL DNAs, revealing that MutSβ-IDL complexes exhibit an overall structural similarity to DNA-bound human MutSα and bacterial MutS-DNA complexes^14^. These structural analyses highlighted several novel features, including well-defined C-terminal dimerization domains (CTDs) of MSH2 and MSH3, which establish nucleotide-binding asymmetry, as well as characteristic DNA bending that varies depending on the size of mis-paired loop. Notably, these studies demonstrated that all four MutSβ-IDL structures lacked nucleotide in the MSH3 ATPase nucleotide-binding pocket while the MSH2 site was occupied by ADP. Moreover, attempts to soak the crystals with ADP or ATP resulted in reduced X-ray diffraction quality of the protein-DNA complexes. In this study, we observed that both the DNA-free MutSβ and its two open forms bound to (CAG)_2_ IDL DNA accommodate ADP at the MSH3 nucleotide-binding pocket. Specifically, the adenosine of ADP is sandwiched between two conserved aromatic residues, Y877 and Y1057, of MSH3. In these two ADP-bound structures of MutSβ, the MSH3 G900 residue in the Walker A motif is repositioned and opened for phosphate binding. In contrast, the MSH3 Y877 residue occupies the adenine-binding site in the canonical form of MutSβ bound to (CAG)_2_ IDL DNA, resulting in the absence of nucleotide at the MSH3 site, which is consistent with the previous crystallographic study (Extended Data Fig. 10a). Conversely, the ATP or ATPγS-bound structures exhibit the closure of nucleotide-binding pockets, which is facilitated by the interplay between the negatively charged γ-phosphate of nucleotides and the positively charged α-helix dipole, along with the involvement of specific serine residues (S723 in MSH2 and S950 in MSH3) located in the opposite subunit.

Recent biochemical and structural studies of human MutSα and its variant, consisting of MSH2 V63E and wild-type MSH6, have proposed that upon initial mismatch binding the MSH2 MMBD moves towards the MSH6 MMBD, forming an interface between the two MMBDs that locks MutSα into a high-affinity state on the mismatched DNA^22^. The MSH2 V63E mutation (a homozygous variant found in the germline of a patient who developed colorectal cancer at approximately 30 years of age) has been shown to affect DNA binding, likely due to a disrupted interface between the two MMBDs. In the canonical form of the wild-type MutSβ bound to (CAG)_2_ DNA, MSH2 V63 appears to play a direct role in MSH3 MMBD binding by forming a hydrophobic cluster with key residues in the interface between the two MMBDs (Extended Data Fig. 10c). The MSH2 V63E variant of MutSβ may therefore alter DNA binding more dramatically than the wild-type protein, decreasing the binding affinity of MutSβ to mismatched DNA. Intriguingly, when bound to the (CAG)_2_ IDL DNA, the wild-type MutSβ was found to adopt a partially open conformation that resembles the low-affinity state of the MutSα MSH2 V63E variant bound to GT-mismatched DNA^22^. Given the absence of the MSH2 MMBD in the cryo-EM map of our novel MutSβ conformational state, as well as an open structure of the MSH2 clamp, we suggest that the newly discovered partially open conformation of DNA-bound MutSβ corresponds to the low-affinity DNA-bound state. Further studies are required to elucidate the precise roles of this partially open state of MutSβ bound to IDL DNA and their functional implications in DNA MMR.

Our cryo-EM analysis of MutSβ bound to homoduplex DNA revealed two different sliding clamp forms and demonstrated conformational dynamics of the MSH2 connector domain. When bound to long plasmid DNAs that are randomly dispersed in vitreous ice on the cryo-EM grid, the MSH2 connector moves away from the DNA without direct interaction, potentially allowing MutSβ to slide along the DNA until it reaches the open end of the DNA. In the case of the relatively short 61 bp homoduplex DNA binding, the MSH2 connector, which is connected to the compact ATP-bound structure, rotates by almost 180° and moves inward toward one end of the central DNA-binding pore, suggesting that the MutSβ sliding clamp may be held near the end of the open-ended DNA.

Based on our observations, we propose the following model for MutSβ (Fig. 7): (Step 1) in its DNA-free form, MutSβ bound by ADP at both ATPase nucleotide-binding sites adopts a relaxed open conformation suitable for DNA binding. (Step 2) Upon DNA scanning, MutSβ initially binds mismatched DNA bases with low affinity through the mismatch-binding domain of MSH3, causing sharp DNA bending through the insertion of the two conserved residues, tyrosine 254 and lysine 255 of MSH3. During this low-affinity binding state, MutSβ still contains ADP at both nucleotide-binding pockets of the MSH2-MSH3 heterodimer. (Step 3) This is followed by the closing of the mismatch-binding domain of MSH2, which enhances DNA binding and results in the high-affinity binding state of MutSβ to DNA.

**Fig. 7.**
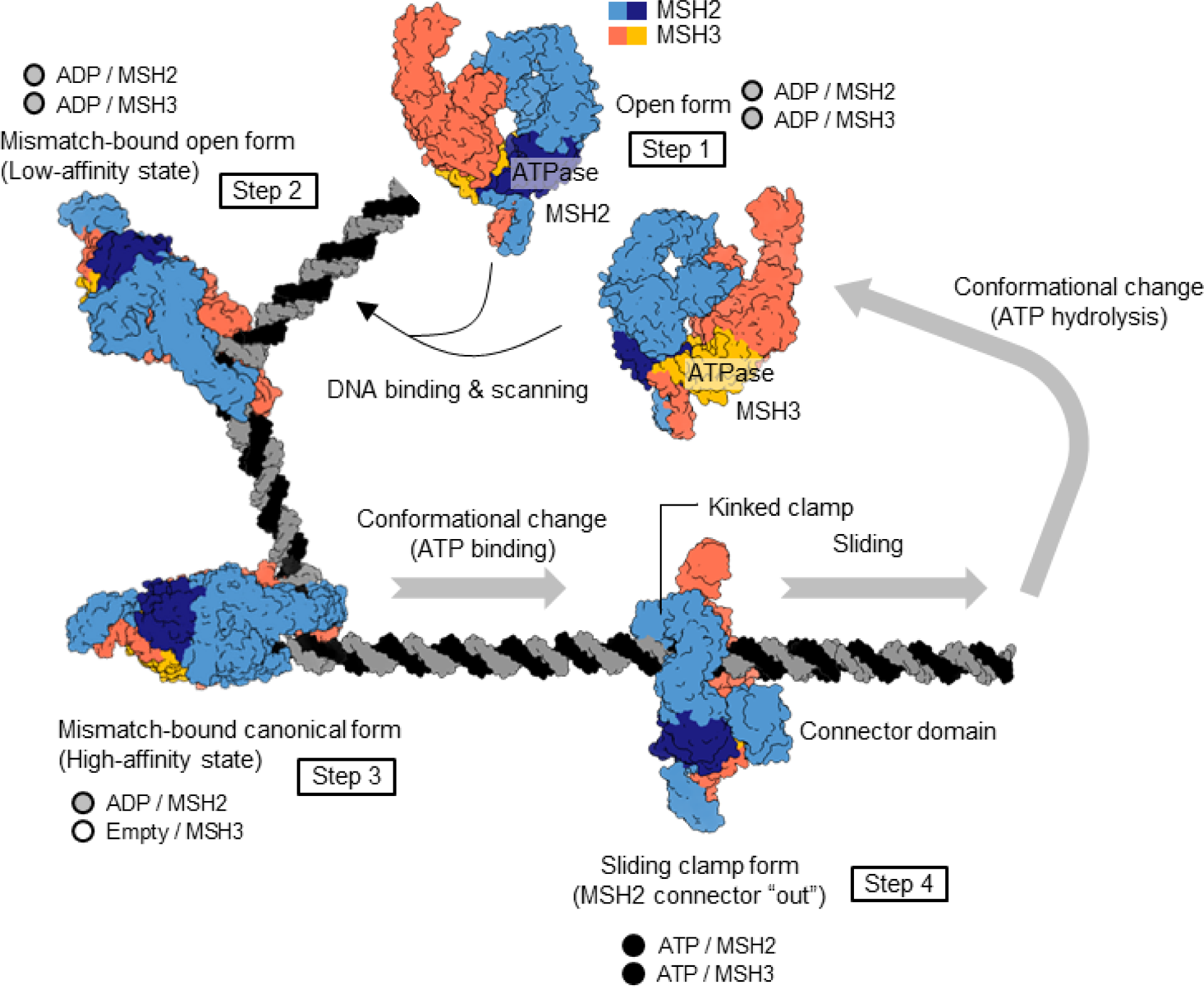
Proposed working model of DNA MMR by MutSβ. Our cryo-EM models depict individual steps of MMR by MutSβ, based on biochemical and structural analyses. The proposed working model further illuminates the dynamic conformational MutSβ cycle for MMR.

Rearrangement of local conformation of key residues forming the MSH3 nucleotide-binding site in the high-affinity state induces ADP release in the MSH3 subunit. (Step 4) MutSβ transitions into a sliding clamp conformation by exchanging ADP for ATP at MSH2 and occupying ATP at MSH3, causing DNA to reposition downward towards the central pore of MutSβ and closing the MSH2 and MSH3 ATPase nucleotide-binding pockets. MutSβ binding to DNA initiates the recruitment of the MutL endonucleases and hence the whole MMR repair pathway. Through the structures described in this study, we can observe snapshots of MutSβ recognizing mismatched DNA and its release from the mismatch in an ATP-dependent manner.

In summary, our structural and biochemical analyses of MutSβ provide key insights into the architecture of MutSβ and its interaction with DNA, highlighting the conformational dynamics that accompany domain rearrangement upon nucleotide and DNA binding and release. Our study contributes not only to the understanding of the dynamic conformational changes of MutSβ and how it initiates the MMR cascade but also provides a comprehensive view of the nucleotide binding pocket to which ATP competitive small molecule ligands are being developed as inhibitors. Additionally, functional and molecular dynamic studies can be employed to identify novel points of intervention for therapies such as allosteric modulators of the ATPase activity and the DNA/protein interface. Improved knowledge of the structure and function of MutSβ will help develop new therapeutic strategies for the treatment for HD and other triplet repeat neurodegenerative disorders.

---

**Extended Data Fig. 1.**
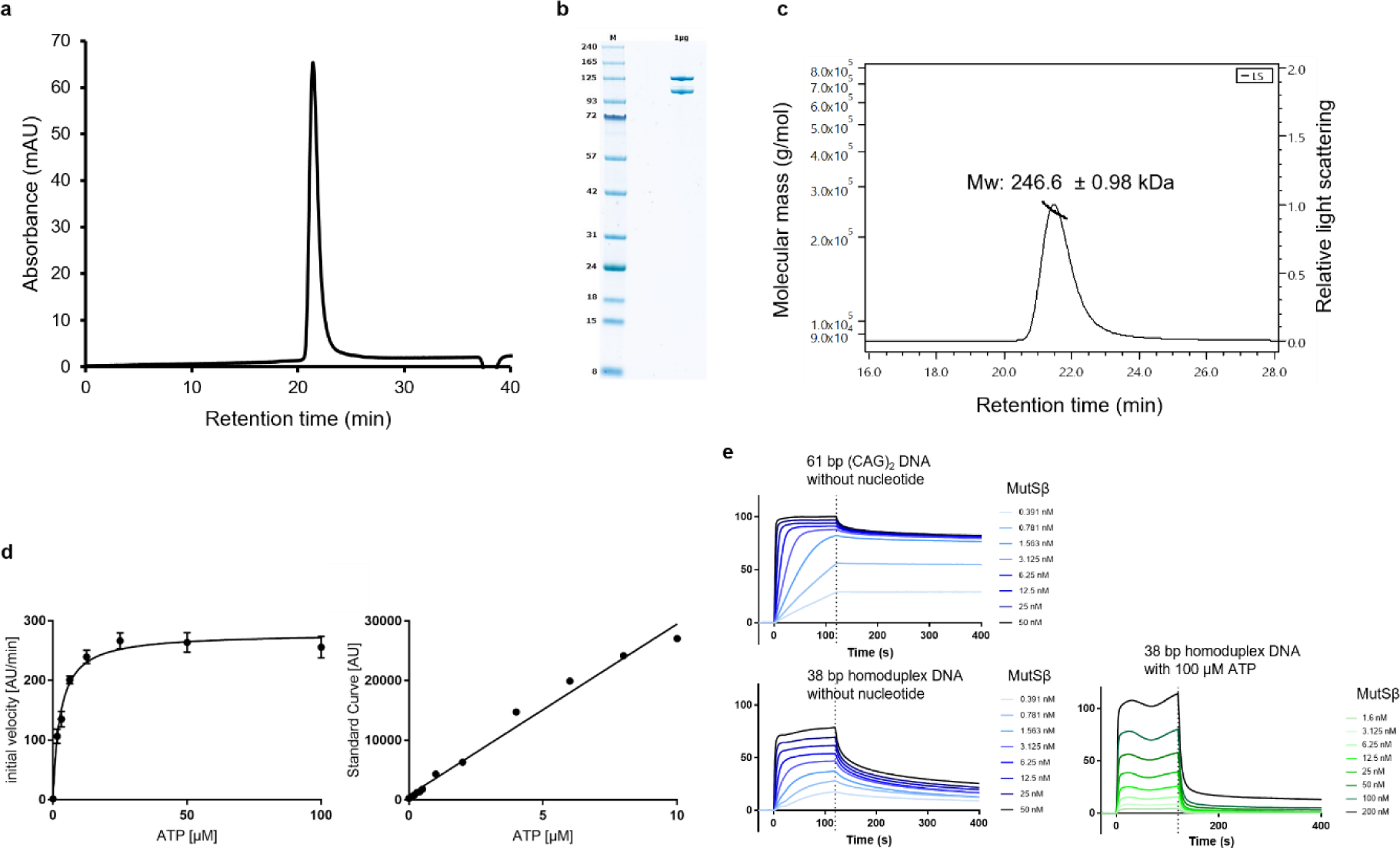
Biochemical analysis of full-length MutSβ. **a,** SEC analysis of purified full-length MutSβ. **b,** SDS-PAGE of purified MutSβ for cryo-EM grid preparation. **c,** SEC-MALS analysis of purified full-length MutSβ. **d,** ATPase activity determination of MutSβ by ADPglo in absence of DNA. Background subtracted initial velocity of ATP dilution series is plotted over ATP concentration (left). Non-linear fit of reaction time yields a V_max_ of 279 AU/min or 0.1 µM/min, converted using a slope of 2,872 AU/[µM] from linear regression (right; V_max_ = 0.12 µM/min; K_M_ = 6.9µM; k_cat_ = 1.22 min^-1^). **e,** DNA binding of MutSβ to immobilized DNA as measured by SPR.

**Extended Data Fig. 2.**
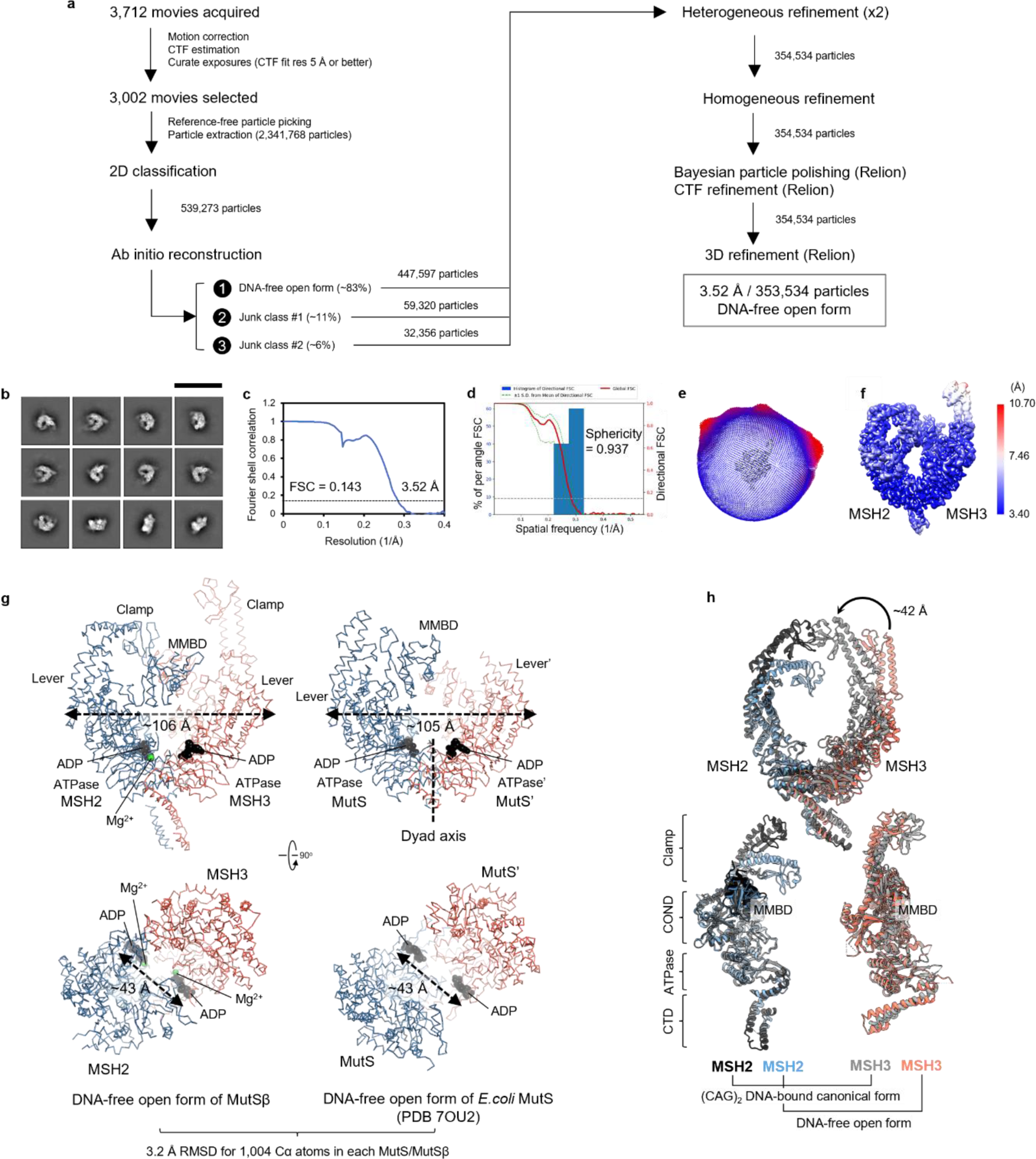
Cryo-EM analysis of DNA-free open form of MutSβ-ADP complex. **a,** Summary of the image processing workflow. **b,** 2D classes of DNA-free open form of MutSβ. The scale bar in white indicates 293 Å. **c,** Gold-standard FSC curve for the density map of DNA-free open form of MutSβ. **d,** 3D FSC^42^ plot for the density map of DNA-free open form of MutSβ. **e,** Heat map showing particle orientation distribution. **f,** Local resolution represented by a heat map on the density contour. **g,** Ribbon diagrams of human MutSβ and *E. coli* MutS (PDB 7OU2)^21^ are shown in their DNA-free open form, based on superposition. **h**, Relative domain movements of the open conformation of MutSβ compared to the canonical mismatch-bound form (PDB 3THZ)^14^ (DNA, MMBD and connector are omitted for clarity).

**Extended Data Fig. 3.**
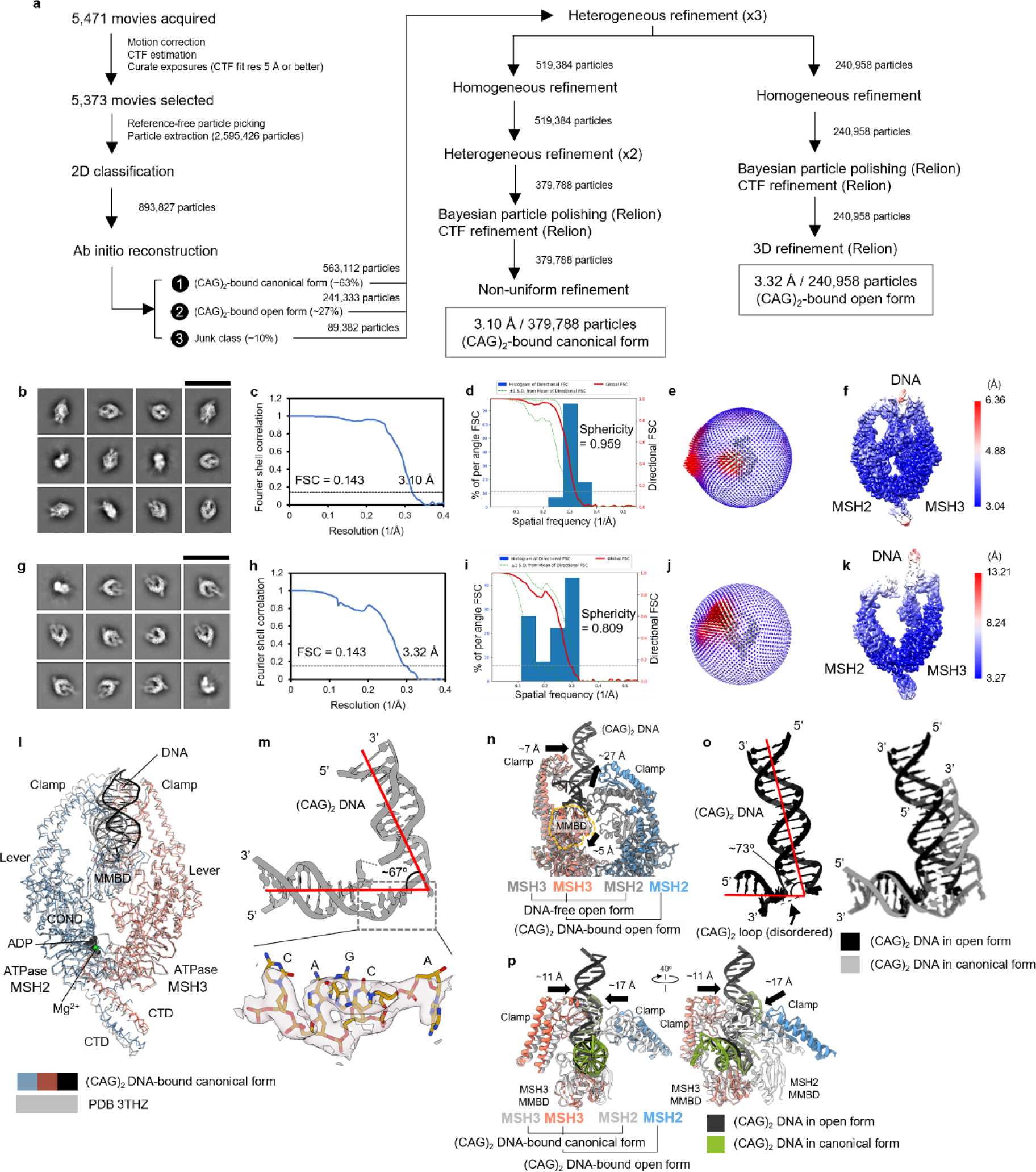
Cryo-EM analysis of (CAG)_2_ DNA-bound canonical and open forms of MutSβ-ADP complex. **a,** Summary of the image processing workflow. **b,** Representative 2D classes of (CAG)_2_ DNA-bound canonical form of MutSβ. The scale bar in white indicates 293 Å. **c,** Gold-standard FSC curve for the density map of (CAG)_2_ DNA-bound canonical form of MutSβ. **d,** 3D FSC^42^ plot for the density map of (CAG)_2_ DNA-bound canonical form of MutSβ. **e,** Heat map showing particle orientation distribution of (CAG)_2_ DNA-bound canonical form. **f,** Local resolution represented by a heat map on the density contour of (CAG)_2_ DNA-bound canonical form. **g**, Representative 2D classes of (CAG)_2_ DNA-bound open form of MutSβ. The scale bar in white indicates 293 Å. **h**, Gold-standard FSC curve for the density map of (CAG)_2_ DNA-bound open form**. i,** 3D FSC^42^ plot for the density map of (CAG)_2_ DNA-bound open form**. j,** Heat map showing particle orientation distribution of (CAG)_2_ DNA-bound open form. **k**, Local resolution represented by a heat map on the density contour of (CAG)_2_ DNA-bound open form. **l,** Ribbon diagram of the mismatch-bound canonical structure of MutSβ, determined by cryo-EM and crystallography (PDB 3THZ)^14^, are shown based on superposition. **m,** The (CAG)_2_ DNA structure in the canonical mismatch-bound state of MutSβ, including a close-up view of mis-paired bases overlaid with electron density (transparent pink) is shown. **n**, Superposition of the DNA-free open conformation of MutSβ with its (CAG)_2_ DNA-bound open state. Substantial domain movements are indicated by arrows. **o**, The (CAG)_2_ DNA structure in the mismatch-bound open state of MutSβ (left) and its overlay with the same DNA in the canonical mismatch-bound state of MutSβ (right) are shown. **p**, Two views of superposition of the two (CAG)_2_ DNA-bound MutSβ cryo-EM structures. The arrows indicate the inward movement of the two clamps in the transition to the canonical high-affinity state.

**Extended Data Fig. 4.**
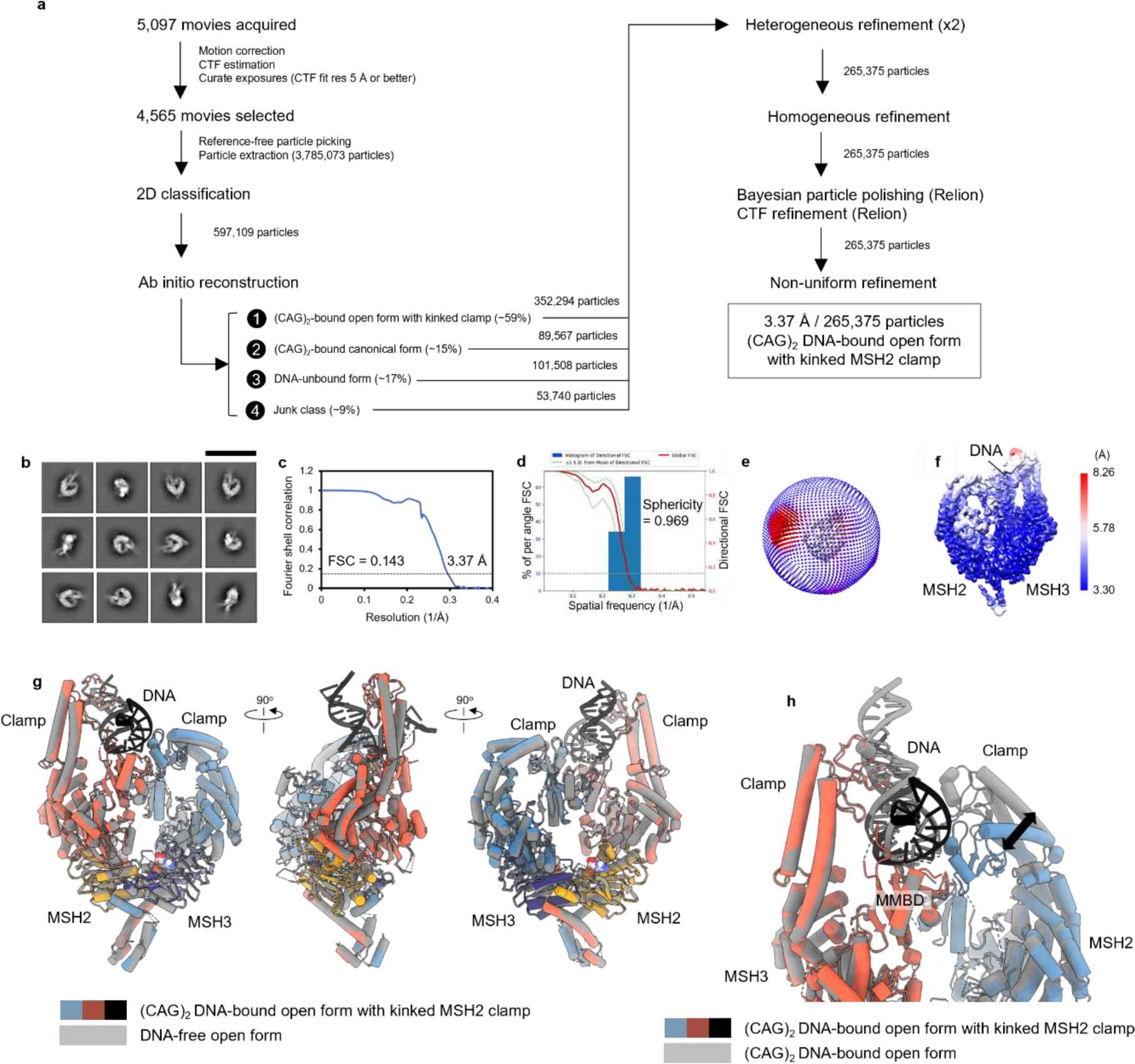
Cryo-EM analysis of (CAG)_2_ DNA-bound open form of MutSβ-ADP complex with kinked MSH2 clamp. **a,** Summary of the image processing workflow. **b,** Representative 2D classes of (CAG)_2_ DNA-bound open form of MutSβ with kinked MSH2 clamp. The scale bar in white indicates 293 Å. **c,** Gold-standard FSC curve for the density map. **d,** 3D FSC^42^ plot for the density map. **e,** Heat map showing particle orientation distribution. **f,** Local resolution represented by a heat map on the density contour. **g,** Three orthogonal views of the (CAG)_2_ DNA-bound open form overlaid with the DNA-free open form of MutSβ are shown based on superposition. **h,** Structural comparison of two cryo-EM structures of MutSβ bound to the same (CAG)_2_ DNA in two different conformations. The black arrow indicates a substantial movement of the MSH2 clamp.

**Extended Data Fig. 5.**
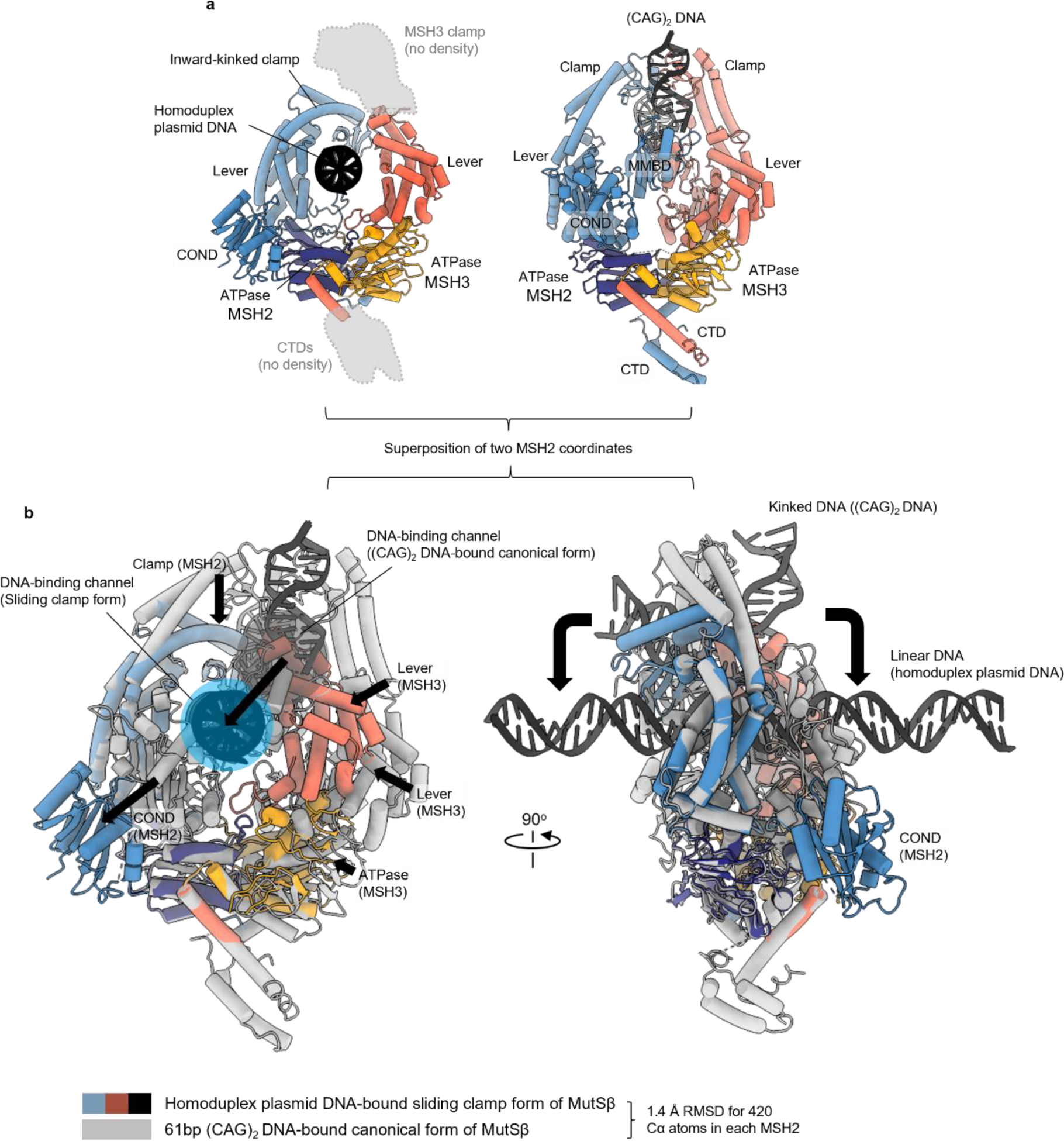
Structural comparison of homoduplex plasmid DNA-bound sliding clamp form of MutSβ with its canonical form bound to (CAG)_2_ DNA. **a,** Ribbon diagrams of human MutSβ in its DNA-bound form are shown for two different conditions: (1) bound to 1.8 kb homoduplex plasmid DNA in the presence of ATP (left) and (2) bound to 61 bp (CAG)_2_ DNA in the absence of added nucleotide (right), based on superposition. **b,** Two orthogonal views of superimposed cryo-EM structures show conformational differences between two structures. The black arrows indicate domain movements of MSH2, MSH3, and DNA towards the homoduplex plasmid DNA-bound sliding clamp state of Mutβ.

**Extended Data Fig. 6.**
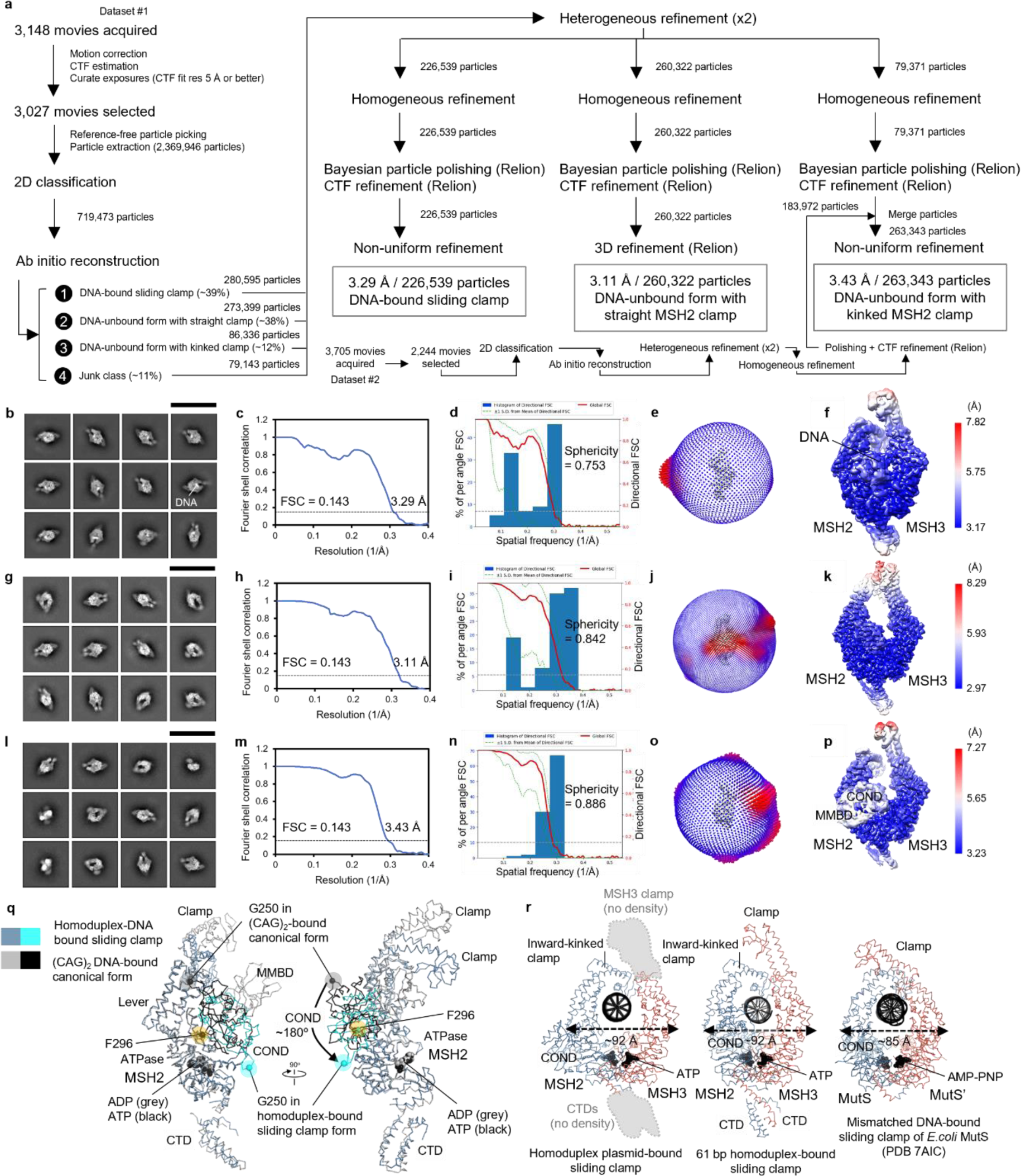
Cryo-EM analysis of 61 bp homoduplex DNA-bound sliding clamp and two DNA-unbound forms of MutSβ-ATP complex. **a,** Summary of the image processing workflow. Representative 2D classes (293 Å scale car in black) (b), gold-standard FSC curve for the density map (c), 3D FSC^42^ plot for the density map (d), heat map showing particle orientation distribution (e), and local resolution represented by a heat map on the density contour (f) of 61 bp homoduplex-bound sliding clamp form of MutSβ. The same corresponding images for DNA-unbound form with straight MSH2 clamp (g-k) and with kinked MSH2 clamp (l-p) are shown in the same order. **q,** Ribbon diagrams of the canonical and sliding clamp forms of MutSβ bound to DNA are shown based on superposition. The MSH2 connector rotates around residue F296 compared to the (CAG)_2_-bound canonical form relative to other domains. The black arrows indicate substantial domain movement of MSH2 connector. **r,** Ribbon diagrams of human MutSβ and *E. coli* MutS (PDB 7AIC)^20^ are shown in their DNA-bound sliding clamp form, based on superposition.

**Extended Data Fig. 7.**
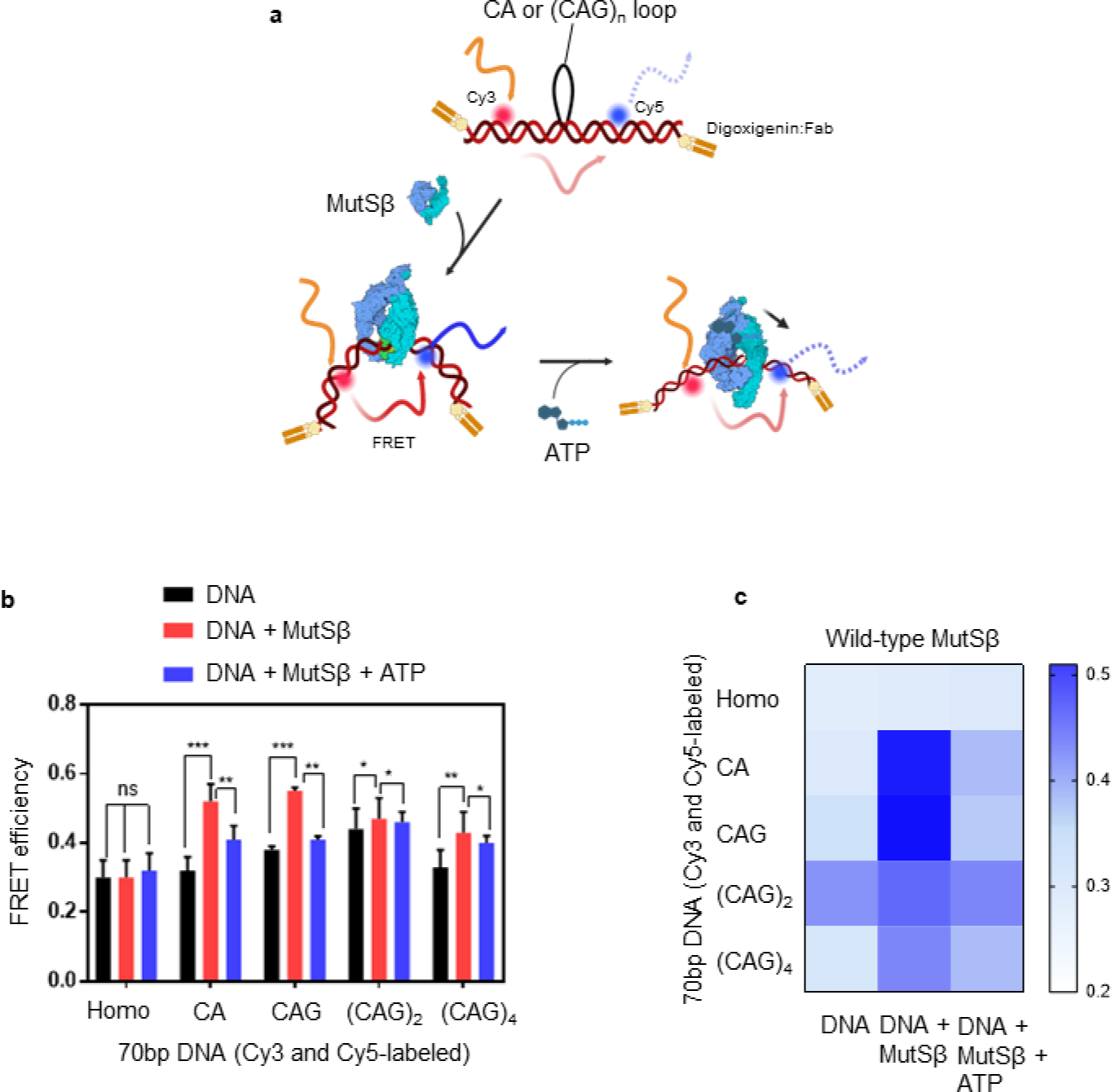
**a,** Schematic representation of DNA substrate to generate a FRET system sensitive to MutSβ DNA mismatch binding due to DNA kinking. 70 bp dsDNA (mismatch shown as black loop and end blocking depicted by antibody binding at 3’ and 5’ ends) was labeled with the donor and acceptor dyes shown as red and blue spheres respectively. FRET signal in the DNA is shown as light colored red and blue arrows. If MutSβ induces kinking in DNA then proximity of the dyes results in more FRET depicted as dark colored arrows. **b**, Comparison of FRET efficiencies of different inserted loops under different conditions is shown in a bar plot. P values are from statistical t-test, n=5 and are depicted as follows: P<0.001 are shown as ***, P<0.005 as ** while P<0.05 are shown as *. **c**, Heat map showing comparison of FRET efficiencies of different mismatched loops for DNA, DNA + MutSβ and DNA + MutSβ + ATP states.

**Extended Data Fig. 8.**
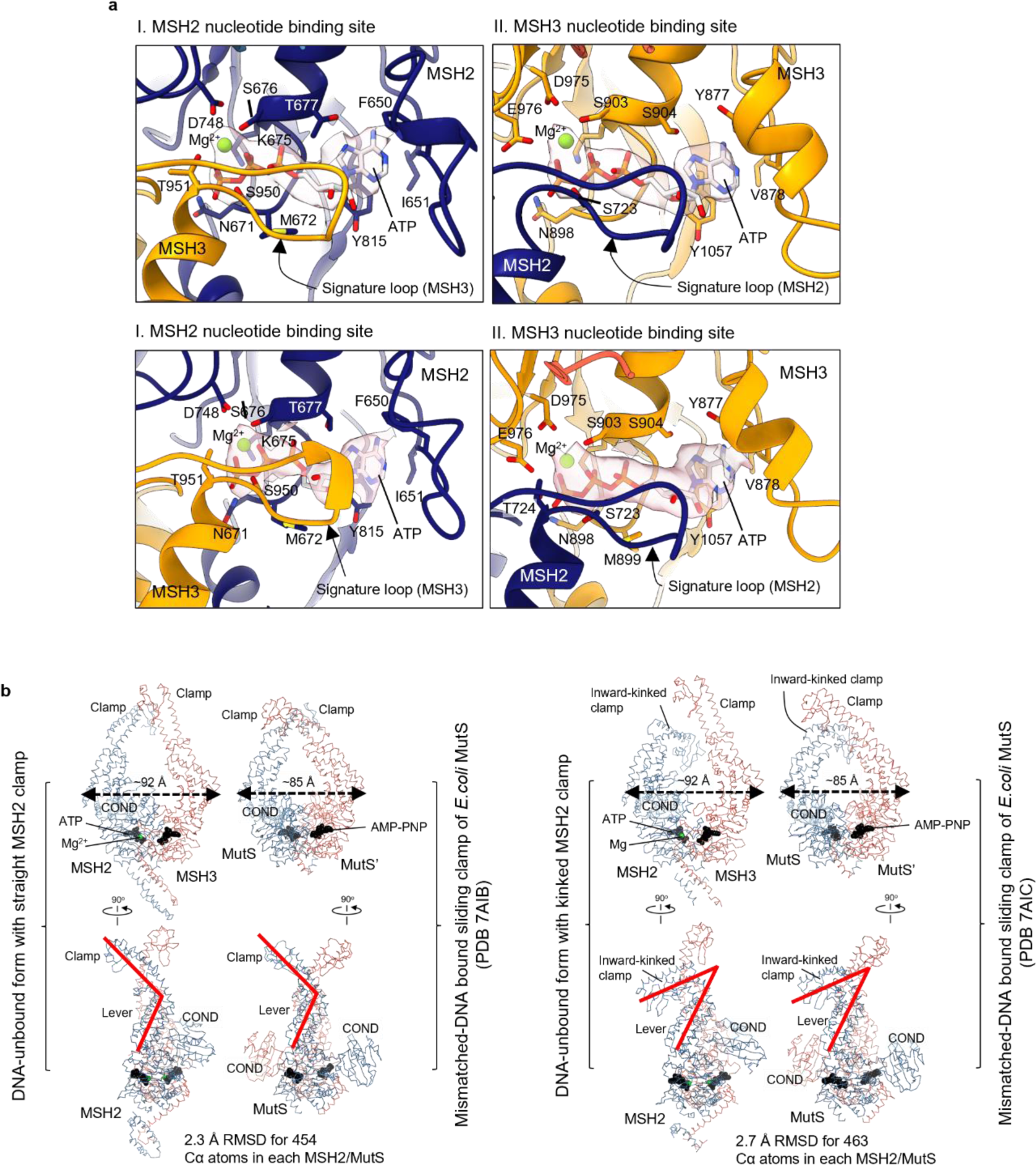
Structural analysis of two DNA-unbound forms of MutSβ-ATP complex. **a,** Close-up views of nucleotide binding pockets in the DNA-unbound MutSβ-ATP complexes: straight MSH2 clamp (top panels) vs. kinked MSH2 clamp (bottom panels). Electron densities for bound ATP-Mg are transparently shown in pink. **b,** Structural comparison of human MutSβ and *E. coli* MutS homodimer (PDB 7AIB and 7AIC; DNA and MutL are omitted for clarity)^18^ are shown in their ATP- or AMP-PNP-bound forms, based on superposition. A sharp kinking of the clamp domain of MSH2 and one subunit of MutS (indicated in red on the right) is shown.

**Extended Data Fig. 9.**
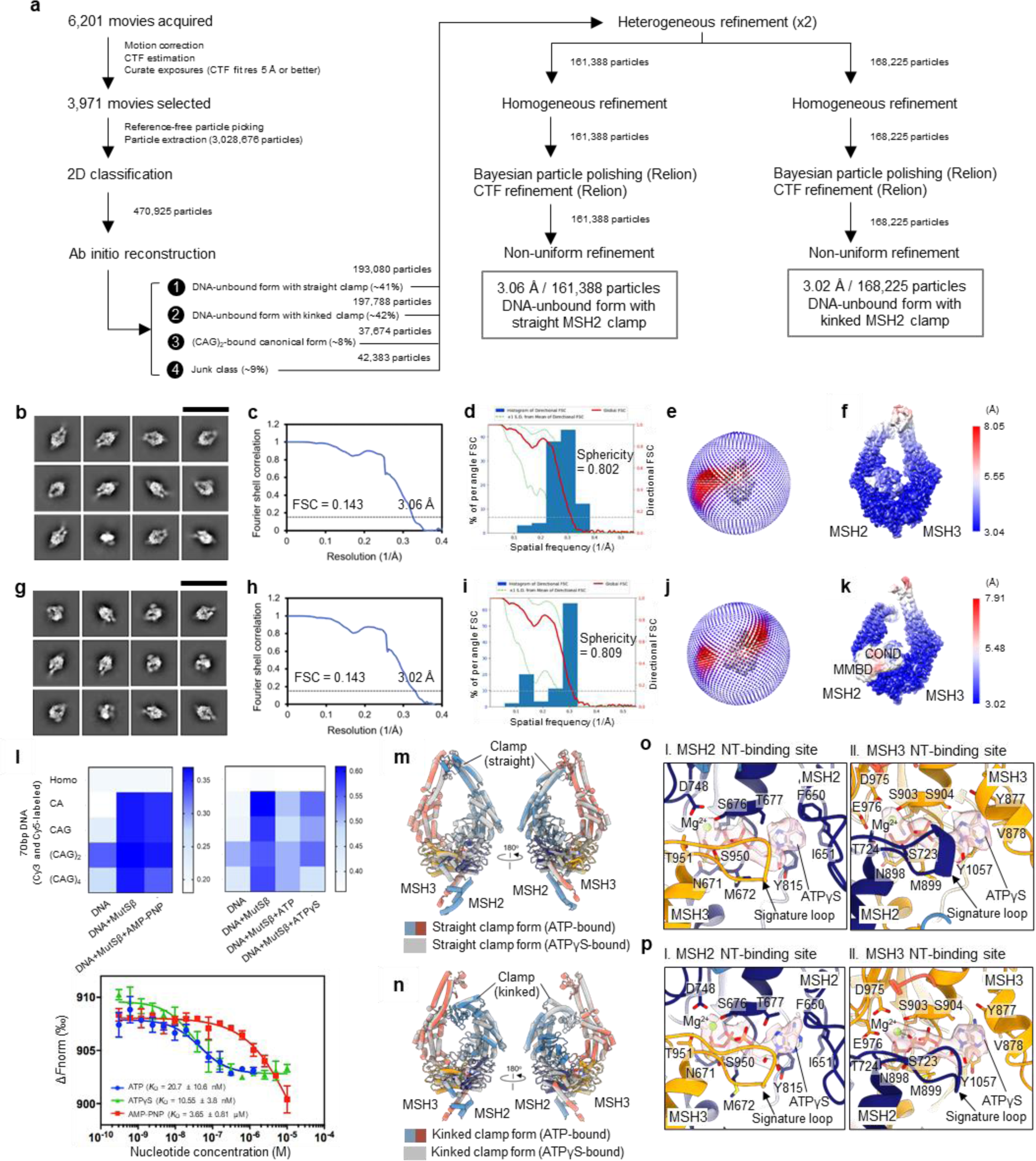
Cryo-EM analysis of DNA-unbound MutSβ-ATPγS complex with straight or kinked MSH2 clamp. **a,** Summary of the image processing workflow. Representative 2D classes (293 Å scale bar in black) (b), gold-standard FSC curve for the density map (c), 3D FSC^42^ plot for the density map (d), heat map showing particle orientation distribution (e), and heat map showing particle orientation distribution (f) of DNA-unbound MutSβ-ATPγS complex with straight MSH2 clamp. The corresponding images for the DNA-unbound MutSβ-ATPγS complex with kinked MSH2 clamp are shown in the same order (g-k). **l**, FRET results displaying nucleotide-dependent signal changes in DNA alone and MutSβ-DNA complexes (top). The binding affinity of three different nucleotide toward the MutSβ-DNA complex was measured from three independent replicates by MST, as shown means ± SD (bottom). **m**, Structural comparison of DNA-unbound forms of MutSβ bound to ATP or ATPγS with straight MSH2 clamp is shown based on superposition. **n**, Structural comparison of DNA-unbound forms of MutSβ bound to ATP or ATPγS with kinked MSH2 clamp is shown based on superposition. Close-up views of nucleotide binding pockets in the DNA-unbound MutSβ-ATPγS complexes: straight MSH2 clamp (o) vs. kinked MSH2 clamp (p). Electron densities for bound ATPγS-Mg are transparently shown in pink.

**Extended Data Fig. 10.**
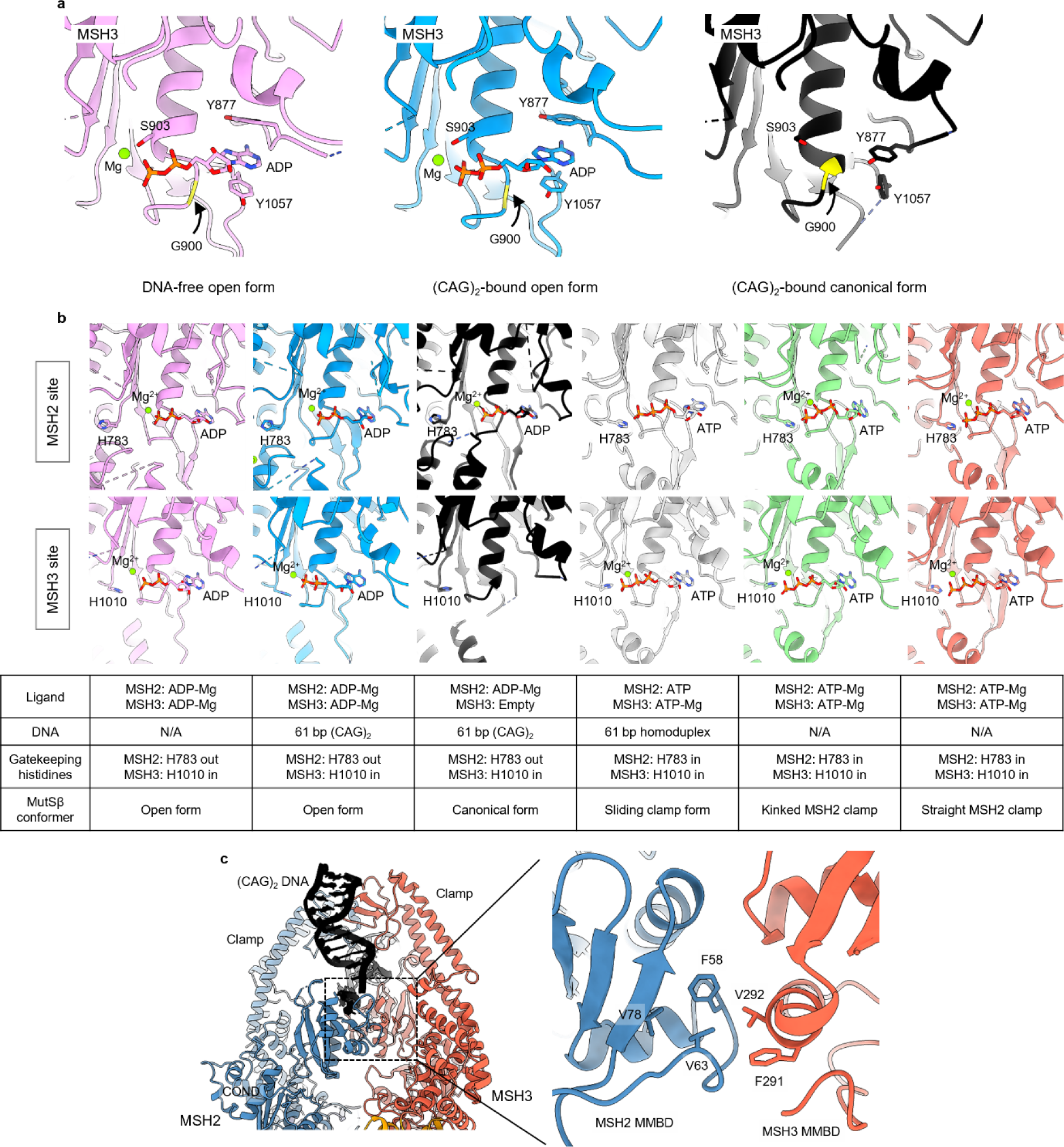
Structural analysis of different conformational states of human MutSβ. **a,** Ribbon diagrams of the MSH3 site of the DNA-free open form of MutSβ (left) and its (CAG)_2_ DNA-bound form (middle and right) are shown based on superposition. The MSH3 G900 residue in the Walker A motif is highlighted in yellow. In the canonical form of MutSβ complexed with (CAG)_2_ DNA, the MSH3 Y877 residue occupies the adenine-binding site, leading to a lack of nucleotide at the MSH3 nucleotide-binding pocket. **b,** Structural comparison of the conformation of gatekeeping histidine residues at the nucleotide-binding pockets of the MSH2 and MSH3 ATPase cores in the six different conformational states of MutSβ. **c,** Hydrophobic interface between MSH2 and MSH3 MMBDs in the (CAG)_2_ DNA-bound canonical form of MutSβ.

**Extended Data Table 1.**
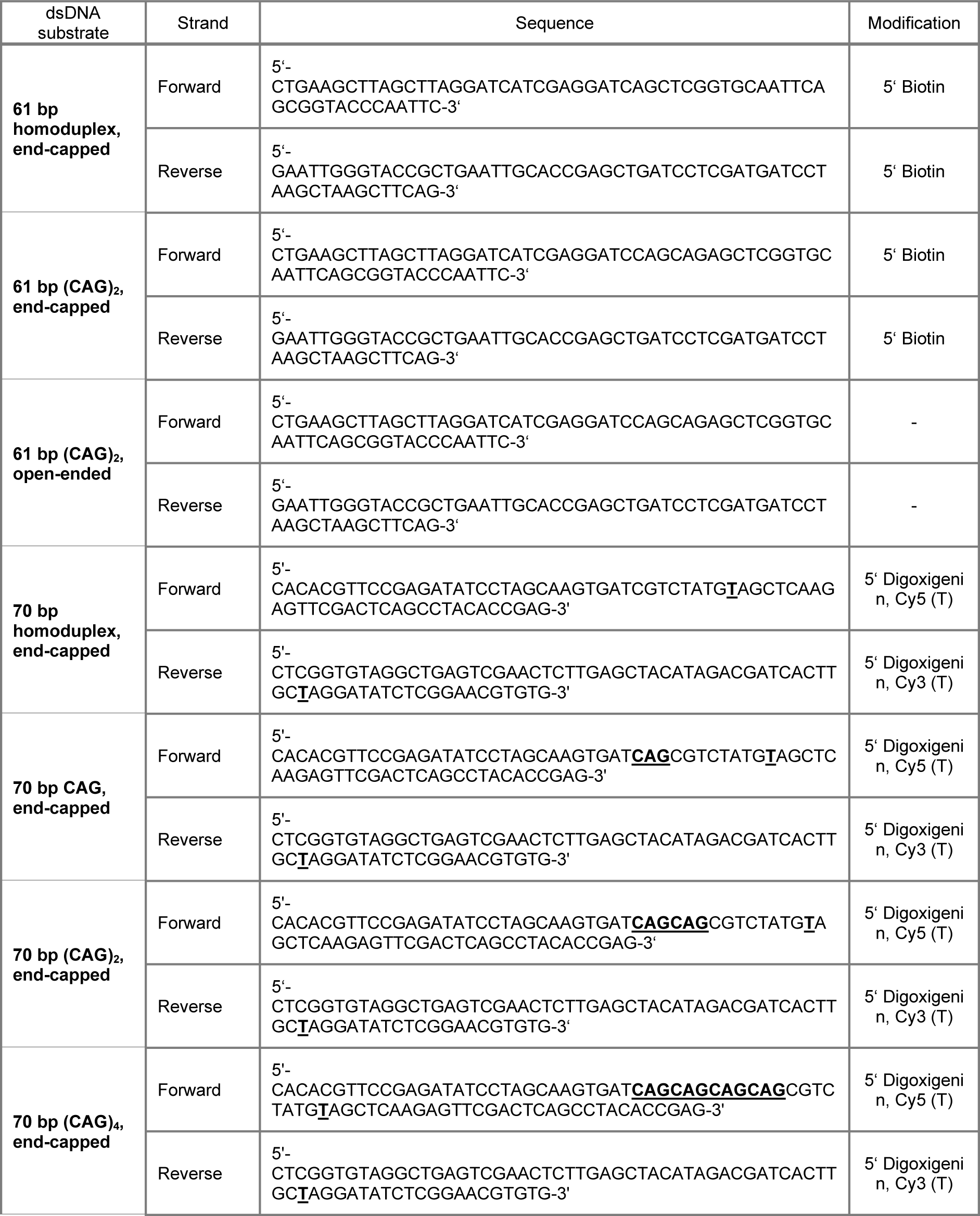
Sequences of dsDNA substrates used in this study.

## Methods

### Plasmid generation

The coding sequences of the human Msh2 (NP_001393570.1) and Msh3 (NP_002430.3) proteins were codon optimized for *S. frugiperda* and synthesized by Genscript (Genscript Biotech, New Jersey, USA) and cloned into a pFastBac-dual vector between the XhoI and NcoI and BamHI and HindIII sites respectively.

### Virus generation & expression

The recombinant bacmid, and baculovirus stocks (P1, P2 & HTVS) were generated according to the manufacturer’s instructions (Invitrogen by Life technologies, https://tools.thermofisher.com/content/sfs/manuals/bactobac_man.pdf). The steps leading to the production of the virus stocks were performed using Sf9 cells and Sf-900™ II medium. The expression of the recombinant proteins was performed using Sf21 cells, with a final expression volume of 10 L of Sf-900™ III medium. The cell pellets were frozen at -80°C until use.

### Protein purification

Pellet from baculovirus infected Sf9 cells expressing the human MutSβ complex were resuspended in lysis buffer (25 mM HEPES pH 8, 1 M NaCl, 10 % glycerol, 1 mM EDTA, 2 mM β-mercaptoethanol) supplemented with protease Inhibitors on ice. Cell lysis was achieved by Turrax homogenization on ice. The lysate was centrifuged at 30,000 g for 10 minutes at 10 °C. The supernatant (clarified lysate) was collected. The clarified lysate was loaded onto a 20 mL DEAE column pre-equilibrated in buffer A (25 mM HEPES pH 8, 1 M NaCl, 10 % glycerol, 1 mM EDTA, 2 mM β-mercaptoethanol) supplemented with protease inhibitors. The bound proteins were eluted from the column by a step of 100 % buffer B (25 mM HEPES pH 7.6, 2 M NaCl, 1 mM EDTA, 2 mM β-mercaptoethanol) supplemented with protease inhibitors. The fractions were analyzed by SDS-PAGE and the fractions containing the MSH2-MSH3 complex were pooled. The pooled fractions from the previous chromatography step were diluted to a final NaCl concentration of 250 mM. The resulting sample was loaded onto a 5 mL Heparin column pre-equilibrated in buffer C (25 mM HEPES pH 7.6, 250 mM NaCl, 1 mM EDTA, 2 mM β-mercaptoethanol) supplemented with protease inhibitors. The bound proteins were eluted from the column by a gradient (0-100 % over 5 column volumes) of buffer D (25 mM HEPES pH 7.6,1 M NaCl, 1 mM EDTA, 2 mM β-mercaptoethanol) supplemented with protease inhibitors. The fractions were analyzed by SDS-PAGE and the fractions containing the MSH2-MSH3 complex were pooled. The pooled fractions from the previous chromatography step were diluted to a final NaCl concentration of 100 mM. The resulting sample was loaded onto a 6 mL Resource Q column pre-equilibrated in buffer E (25 mM HEPES pH 7.6, 100 mM NaCl, 1 mM EDTA, 2 mM β-mercaptoethanol) supplemented with protease inhibitors. The bound proteins were eluted from the column by a gradient (0-100 % over 5 column volumes) of buffer D (25 mM HEPES pH 7.6, 1 M NaCl, 1 mM EDTA, 2 mM β-mercaptoethanol) supplemented with protease inhibitors. The fractions were analyzed by SDS-PAGE and the fractions containing the MSH2-MSH3 complex were pooled. The pooled fractions from the previous chromatography step were diluted to a final NaCl concentration of 100 mM. The resulting sample was loaded onto a 1 mL Mono Q column pre-equilibrated in buffer E (25 mM HEPES pH 7.6, 100 mM NaCl, 1 mM EDTA, 2 mM β-mercaptoethanol) supplemented with protease inhibitors. The bound proteins were eluted from the column by a gradient (0-100% over 5 column volumes) of buffer D (25 mM HEPES pH 7.6, 1 M NaCl, 1 mM EDTA, 2 mM β-mercaptoethanol) supplemented with protease inhibitors. The fractions were analyzed by SDS-PAGE and the fractions containing the MSH2-MSH3 complex were pooled. The pool from the previous purification step was concentrated to a volume of 10 mL and loaded onto a S200 26/60 column pre-equilibrated in buffer G (25 mM HEPES pH 7.5, 100 mM KCl, 5 mM MgCl_2_, 0.1 mM EDTA, 1 mM TCEP, 5% glycerol). The fractions were analyzed by SDS-PAGE and the fractions containing the MSH2-MSH3 complex were pooled. The fractions pooled from the size-exclusion chromatography step were concentrated using an Amicon concentrator (MWCO 50 kDa) to a final concentration of 13 mg/mL.

### Generation of dsDNA substrates

The 61 bp single-stranded DNAs were synthesized by Metabion (Metabion International AG, Planegg, Germany), the sequences can be found in table 2. Equimolar amounts of each ssDNA molecule were mixed to a final concentration of 500 µM each in TE Buffer (10 mM Tris-HCl pH 8, 1 mM EDTA). Annealing was performed by heating the DNA sample to 95 °C for 2 minutes and subsequently cooling from 95 °C to 20 °C with a 1 °C/min ramp (Labcycler, SensoQuest GmbH). In the case of the biotinylated dsDNA substrates, pre-saturation of Streptavidin with Biotin was achieved by mixing Streptavidin (AnaSpec Inc.) to Biotin (Merck Millipore) in a molar ratio of 1:3.5 at 500 µM Streptavidin and 1,750 µM Biotin in 100 mM Tris-HCl pH 8.0, 100 mM NaCl, 5 mM MgCl_2_, incubating for 3 h at ambient temperature. Subsequently, biotinylated dsDNA samples were end-blocked by addition of 1 eq. of 61 bp 5’-biotinylated homoduplex dsDNA to 16 eq. of pre-saturated Streptavidin at a total volume of 36 µL, incubated over night at 4 °C. This corresponds to a final DNA concentration of 27.78 µM with a final Streptavidin concentration of 444.44 µM. Purification of the Strep-capped DNA was achieved by loading the sample mixture on a Pierce™ Strong Anion Exchange Spin Column and washing with 400 µL of 50 mM Tris pH 8.0, 5 mM MgCl_2_ with increasing NaCl concentration, starting from 200 mM NaCl. Spin columns were equilibrated and handled according to manufacturer’s instructions. Columns were conditioned twice using 400 µL of 50 mM Tris-HCl pH 8.0, 200 mM NaCl, 5 mM MgCl_2_. Samples were pre-washed by addition of 50 mM Tris-HCl pH 8.0, 5 mM MgCl_2_ with increased NaCl concentrations from 200 to 600 mM in 100 mM increments. From 600 mM to 800 mM, the sodium chloride concentration was increased in 25 mM increments to elute the Streptavidin-end-blocked DNA from the column. Absorbance spectra of elution fractions were measured and pooled according to an absorbance ratio of A260/A280 of 1.00 to 1.65. The linear plasmid substrate was generated by PCR from a pET16b vector to generate a 1,783 bp fragment comprising the ORI and AmpR gene and promoter of the pET16b vector.

### SEC-MALS analysis

For the determination of the molecular mass of MutSβ, SEC-MALS was applied. Therefore, an Ultimate 3000 HPLC system (Thermo Fisher Scientific) equipped with a degasser, a quaternary pump, an autosampler (kept at 10 °C) and a VWD detector was coupled to a miniDawn TREOS II 3-angle light-scattering detector and an Optilab RI detector (Wyatt Technology). The HPLC system was controlled by Chromeleon 7.3.1 (Thermo Fisher Scientific) and the MALS as well as the RI detector with Astra 7.3.2.21 (Wyatt Technology). For the analysis the protein solution (100 µL, 1 mg/mL) was injected on a Superdex 200 Increase 10/300 GL (30 cm × 10 mm, 8.6 μm particle size) SEC column. As a mobile phase the following buffer was used: 25 mM HEPES, 100 mM KCl, 5 mM MgCl_2_, 1 mM TCEP, 0.5 mM EDTA, pH7.5. All measurements were conducted at room temperature. Data analysis was performed using Chromeleon 7.3.1 (HPLC-UV data) and Astra 7.3.2.21 (MALS data).

### ATPase activity assay

ATPase activity was determined by ADP-Glo Kinase Assay Kit (Promega, V9102), following the manufacturer’s protocol. Initially, a 384-well plate (Corning® 384-well Low Volume White, product number 4513) was prepared for reaction stop. All wells were prefilled with 4 µl ADP-GloTM reagent. Storage at room temperature. Then, a master mix plate was prepared for the ATPase reaction. A serial two-fold dilution of ATP was prepared in Eppendorf tubes (maximum ATP conc. = 200 µM, 2-fold dilution series, eight titration points). 30 µl of ATP dilutions were filled in the respective columns of Row A and Row B of a second 384-well plate (Greiner product number 781270) with columns 22-24 = maximum ATP concentration, columns 1-3 = 0 mM ATP. MutSβ samples were diluted in reaction buffer to a concentration of 200 nM in an Eppendorf tube. The reaction was started by transferring 30 µl of MutSβ samples to the preplated ATP dilution series in Row A using a Multipette (Eppendorf), hereafter referred to as “reaction mix”. For the background control section, 30 µl buffer were added to row B, hereafter name “background control mix”. Prepared plates were thoroughly mixed using MixMate (Eppendorf), followed by incubation at 37 °C. Final enzyme concentration was 100 nM, with a maximum ATP concentration of 100 µM. For the 0 min reaction time point, 4 µl ATP from the dilution series prepared for the master mix plate (Eppendorf tubes with maximum ATP conc. = 200 µM) were added to row A and row I on reaction stop plate. Transfer of 4 µl of prediluted 200 nM MutSβ protein solution to row A or of buffer to row I, followed by thorough mixing using MixMate (Eppendorf). For all further reaction time points, 8 µl of ATP-ATPase reaction mix were transferred from row A on master mix plate to respective row (B to H) on reaction stop plate using a Matrix^TM^ Multichannel Pipette (Thermo Fisher). Additionally, transfer of 8 µl background control mix from row B of master mix plate to respective row (J to P) on reaction stop plate, followed by thorough mixing using MixMate (Eppendorf). Samples were taken at 15, 30, 45, and 60 min timepoints. Subsequently, the detection reaction was performed by 40 min incubation at room temperature with ADP-Glo reagent, followed by addition of 6 µl kinase detection reagent. Plates were thoroughly mixed using MixMate (Eppendorf). After another incubation for 40 min at room temperature, the readout of luminescence signal was recorded using a LUMIstar Omega (BMG Labtech). For each run, a standard curve was created that represents the luminescence signal corresponding to the ATP converted to ADP. Therefore, ATP and ADP stock solutions that was provided with the Assay kit were mixed in different ratios to represent 100 % conversion at 10 µM ADP/0 µM ATP and 0 % conversion at 0 µM ATP/10 µM ADP, as well as linear combinations of both ADP and ATP in between. Collected data was first corrected for background control signal by subtraction from the reaction mix signal at the respective time point. The background subtracted ADP-Glo Signal was plotted versus time for each concentration of ATP and fitted to a linear regression model. Linearity was granted for R-squared ≥ 0.95. At low ATP concentrations, this criterion was not met for long reaction time points as all ATP was consumed such that a plateau was reached. These data points were excluded from data evaluation. The slope represents the reaction velocity in luminescence counts/min for each ATP concentration. The slope was plotted versus ATP concentration and fitted to the Michaelis Menten Equation (Equation 1).

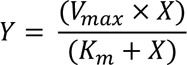

Equation 1: K_m_ represents the ATP concentration at half maximal reaction velocity. V_max_ represents the maximum reaction velocity in luminescence counts/min. The standard curve was fitted to a linear regression and the slope in luminescence counts/µM was used to convert the V_max_ value to µM/min. The turnover number k_cat_ in min^-1^ was determined by dividing the V_max_ value by the enzyme concentration.

### Surface plasmon resonance

SPR binding experiments were performed on a Biacore 8K instrument (Cytiva) at T = 25°C. Biotinylated DNA oligos were annealed to dsDNA in solution and subsequently immobilized on a streptavidin-coated SA chip Series S (Cytiva). Immobilization was performed using a DNA concentration of 1 nM. Immobilization levels were kept between 10 and 30 RUs, depending on DNA oligo length. Running buffer was 25 mM HEPES, pH 7.5, 150 mM KCl, 5 mM MgCl_2_, 1 mM DTT, 0.05% Tween and 0.05% BSA. Samples were kept at 20°C until injection. Binding studies with MutSβ are performed using 2-fold dilution series. Experimental parameters for kinetics experiments were chosen as follows: MutSβ in presence of ATP [1.56 – 200 nM] in absence of ATP [0.39 – 50 nM] was injected for 120 s and dissociation was monitored for 300 s at a flow rate of 30 µl/min. After each MutSβ injection, the chip surface was regenerated with a solution of 0.5% SDS. For assessment of ATP effects, running buffer was supplemented with 0.1 mM ATP. Kinetics data from SPR measurements were analyzed using Biacore Insight Evaluation software v4.0.8.20368. Dissociation constants (KD_eq_) were obtained from steady state affinity global fitting. Information on kinetic rate constants and residence times were extracted using a 1:1 binding model. During the experiments, we checked that calculated maximum binding (theoretical R_max_) was in line with immobilized oligonucleotide. Graphs were created using GraphPad Prism v7.05.

### FRET assay

70 bp DNA with different mismatched loops (homoduplex, CA, CAG_1_, CAG_2_, CAG_4_) were used in the present study (Extended Data Table 1). The forward sequence was labeled with the Cy-5 and the reverse sequence with Cy-3 fluorophores at the position X (dT) resulting in an 5’-X-9bp-(CAG)_n_-8bp-X-3’ arrangement upon complementation. Appropriate donor only and acceptor only controls having only one of the dsDNA strands labeled with either donor or acceptor dye were also produced for each DNA substrate type (homo or mismatched loops). The single labeled DNA strands were annealed in Tris-EDTA buffer, pH 7.5 (heating at 95°C and cooling down with a gradient of 1°C/min to RT). Annealed dsDNA was stored at -20°C at a concentration of 50 µM. The dsDNA was end blocked at a concentration 1000 nM with Anti-Digoxigenin Fab Ab fragments (Roche) for 15 min at RT in the assay buffer: 50 mM Hepes pH 7.0, 200 mM NaCl, 1 mM MgCl_2_, 1 mM DTT, 0.05% Tween-20. Labeled dsDNA was used at 5 nM in the assay. For samples with the protein, labeled DNA was incubated with 100 nM of MutSβ for 30 min at RT before FRET measurements. ATP was used at a final concentration of 500 µM in ATP containing samples. Assay was performed in a 384 well plate with 6 µl total assay volume. Fluorescence in the donor (545/590 nm) and acceptor channel (545/680 nm) were measured after donor excitation at 545 nm using Pherastar BMG plate reader. The corrected FRET efficiency E, taking into account respective errors in the acceptor channel, was calculated using the following formula^43^:

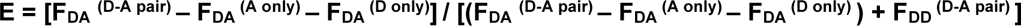

Where F_DA_ represents donor channel excitation/acceptor channel emission and F_DD_ represents donor channel excitation/donor channel emission respectively. D-A pair means double labeled species while D only and A only means donor only and acceptor only labeled dsDNA respectively. Correction factors are given by F_DA_ ^(A only)^ which takes into account direct excitation of acceptor by the donor wavelength, 545 nm and F_DA_ ^(D only)^ corrects for the cross talk of donor in the acceptor channel.

### Cryo-EM sample preparation

Four microliters of full-length wild-type MutSβ at 3 µM were mixed with DNA (3.5 µM of a 61 bp open-ended (CAG)_2_ DNA, 50 nM of linearized plasmid DNA, or 3.5 µM of a 61 bp end-blocked homoduplex DNA). In both DNA-bound sliding clamps and DNA-unbound straight and kinked MSH2 clamp forms, 1 mM ATP was added to the protein-DNA mixtures for the formation of MutSβ-ATP complexes. For the DNA-unbound form of MutSβ-ATPγS with straight or kinked MSH2 clamp, the MutSβ bound to a 61 bp open-ended (CAG)_2_ DNA was mixed with 1 mM ATPγS. To enhance the percentage of particles belonging to the DNA-unbound form of MutSβ-ATP complex with the kinked MSH2 clamp conformer, we preincubated MutSβ with a 61 bp open-ended (CAG)_2_ DNA, followed by the addition of 1 mM ATP. To obtain the (CAG)_2_ DNA-bound MutSβ with the kinked MSH2 clamp, we preincubated MutSβ and a 61 bp open-ended (CAG)_2_ DNA, followed by the addition of 1 mM AMP-PNP. No DNA or nucleotide were added to the protein for the DNA-free open form of MutSβ. All cryo-EM samples mixed with 0.006 % Tween-20 were applied onto a glow-discharged R1.2/1.3 cooper or gold 300-mesh holey carbon grid (Quantifoil) and were immediately frozen in a liquid ethane-propane mixture using a Vitrobot Mark IV (Thermo Fisher Scientific) with the settings at 4 °C, 95 % humidity, and 3-6 seconds of blot time.

### Cryo-EM data processing and image processing

All cryo-EM data were acquired at the cryo-EM facility at Proteros Biostructures GmbH (Table 1). The data were acquired on a Glacios transmission electron microscope (Thermo Fisher Scientific) operated at 200 keV and equipped with a Falcon IV electron detector. The data were collected at a nominal dose of 55.95-59.93 e^-^/Å^2^ with 40 frames per movie and a pixel size of 0.9142 Å. The target defocus range was between 0.7 and 2.5 µm. A total of 3,712, 5,471, 5,471, 5,097, 571, 3,148, 3,148, 3,148 (data #1)/3,705 (data #2), 6,201, and 6,201 movies were collected for the DNA-free open form, 61 bp (CAG)_2_ DNA-bound open form, 61 bp (CAG)_2_ DNA-bound canonical form, 61 bp (CAG)_2_ DNA-bound open form with kinked MSH2 clamp, ∼1.8 kb linearized homoduplex plasmid-bound sliding clamp form, 61 bp homoduplex DNA-bound sliding clamp form, DNA-unbound form of the bound ATP with straight MSH2 clamp, DNA-unbound form of the bound ATP with kinked MSH2 clamp, DNA-unbound kinked MSH2 clamp, DNA-unbound form of the bound ATPγS with straight MSH2 clamp, and DNA-unbound form of the bound ATPγS with kinked MSH2 clamp, respectively. All cryo-EM data were recorded using EPU (Thermo Fisher Scientific). The individual dataset was imported into Relion^34^, and the movie frames were aligned using Relion’s MOTIONCOR implementation. The motion-corrected micrographs were then fed into cryoSPARC^35^, and the CTF (Contrast Transfer Function) was estimated using CTFFIND. Micrographs that were unsuitable for image analysis (e.g. due to significant drift or heavy contamination with crystalline ice) were removed by manual inspection. Particles were picked with cryoSPARC’s reference-free blob picker from the selected micrographs. An initial set of particles was extracted, and multiple rounds of 2D classification were performed to clean up the individual dataset. Ab initio model generation was carried out using cryoSPARC, followed by hetero refinement. A subset of particles attributed to the best class was subjected to homo refinement using cryoSPARC, followed by Bayesian polishing and CTF refinement using Relion. The final set of particles was refined using either cryoSPARC’s non-uniform refinement or Relion’s 3D refinement.

### Model building and refinement

To build atomic structures of nine different conformations of MutSβ, the crystal structure of the human MutSβ-DNA complex (PDB 3THZ)^14^ or the alternative template structures (Table 1) was fitted into the refined 3D reconstruction maps using UCSF (University of California at San Francisco) Chimera^36^. The structures were then manually rebuilt in COOT^37^ to fit the densities, guided mainly by bulky side-chain residues for sequence assignment. The final atomic structures were refined using CCPEM Refmac^38^ and validated using MolProbity^39^. Structure analysis was performed in COOT^37^, and figures were prepared using PyMOL^40^ and ChimeraX^41^.

To generate a low-resolution model of the homoduplex plasmid DNA-bound sliding clamp form of MutSβ, the 3.43 Å cryo-EM structure of the DNA-unbound kinked MSH2 clamp structure was rigid-body fitted into the map. The MSH2 connector was individually fitted into the density. The MSH3 clamp and two CTD domains, which are not defined in the low-resolution map, were removed. Next, the ideal B-form dsDNA was generated using COOT^37^ and fitted into the density of the central channel of MutSβ.

## Data availability

Cryo-EM maps have been deposited in the Electron Microscopy Data Bank (EMDB) under accession numbers EMD-16969 (DNA-free open form), EMD-16971 (mismatch-bound open form), EMD-19605 (mismatch-bound open form with kinked MSH2 clamp), EMD-16964 (canonical mismatch-bound form), EMD-16972 (homoduplex-bound sliding clamp form), EMD-16975 (plasmid-bound sliding clamp form), EMD-16973 (DNA-unbound MutSβ-ATP with kinked MSH2 clamp), EMD-16974 (DNA-unbound MutSβ-ATP with straight MSH2 clamp), EMD-19607 (DNA-unbound MutSβ-ATPγS with kinked MSH2 clamp), and EMD-19606 (DNA-unbound MutSβ-ATPγS with straight MSH2 clamp). Model coordinates have been deposited in the Protein Data Bank (PDB) under accession codes 8OM5 (open form), 8OM9 (mismatch-bound open form), 8RZ7 (mismatch-bound open form with kinked MSH2 clamp), 8OLX (canonical mismatch-bound form), 8OMA (homoduplex-bound sliding clamp form), 8OMO (DNA-unbound MutSβ-ATP with kinked MSH2 clamp), 8OMQ (DNA-unbound MutSβ-ATP with straight MSH2 clamp), 8RZ9 (DNA-unbound MutSβ-ATPγS with kinked MSH2 clamp), and 8RZ8 (DNA-unbound MutSβ-ATPγS with straight MSH2 clamp).

## Acknowledgements

We thank Simon Noble, Titia Sixma, and Paul Modrich for comments.

## Author contributions

M.T., J.-H.L., H.W., R.R.I., N.P., D.P.F., T.S., M.F., T.F.V., and B.C.P. conceived the study. G.T. cloned and purified all described proteins and prepared linearized plasmid DNA. H.D. prepared end-blocked dsDNA samples and performed all biophysical experiments in this study. A.S. performed ADPglo experiments. H.D. and V.D. performed FRET assay and analysis. J.-H.L. carried out grid freezing, data collection and data processing of all described structures in this study. All authors were involved in analyzing data and manuscript writing.

## Competing interests

This work was supported by the nonprofit CHDI Foundation Inc. CHDI Foundation is a nonprofit biomedical research organization exclusively dedicated to collaboratively developing therapeutics that substantially improve the lives of those affected by Huntington’s disease. CHDI Foundation conducts research in a number of different ways; for the purposes of this manuscript, all research was conceptualized, planned, and directed by all authors and conducted at Proteros Biostructures GmbH.

**Correspondence and requests for materials** should be addressed to Brinda Prasad.

